# Insights into single hiPSC-derived cardiomyocyte phenotypes and maturation using ConTraX, an efficient pipeline for tracking contractile dynamics

**DOI:** 10.1101/2021.03.18.436014

**Authors:** Gaspard Pardon, Henry Lewis, Alison S. Vander Roest, Erica A. Castillo, Robin Wilson, Aleksandra K. Denisin, Cheavar A. Blair, Foster Birnbaum, Colin Holbrook, Kassie Koleckar, Alex C-Y Chang, Helen M. Blau, Beth L. Pruitt

**Affiliations:** Departments of Mechanical Engineering and Bioengineering, Stanford University, School of Engineering and School of Medicine, Stanford, CA, USA; Baxter Laboratory for Stem Cell Biology, Department of Microbiology and Immunology, Institute for Stem Cell Biology and Regenerative Medicine, Stanford University School of Medicine, Stanford, CA, USA; Stanford Cardiovascular Institute, Stanford University School of Medicine, Stanford, CA, USA; Department of Pediatrics (Cardiology), Stanford University School of Medicine, Stanford, CA, USA; Shanghai Institute of Precision Medicine and Department of Cardiology, Ninth People’s Hospital, Shanghai Jiao Tong University School of Medicine, Shanghai 200125, China; Division of Cardiovascular Medicine, Stanford University School of Medicine, Stanford, CA, USA; Department of Mechanical Engineering and Biomolecular Science and Engineering Program, Center for Bioengineering, University of California, Santa Barbara, CA, USA

## Abstract

Cardiomyocytes derived from human induced pluripotent stem cells (hiPSC-CMs) are powerful *in-vitro* models to study the mechanisms underlying cardiomyopathies and cardiotoxicity. To understand how cellular mechanisms affect the heart, it is crucial to quantify the contractile function in single hiPSC-CMs over time, however, such measurements remain demanding and low-throughput, and are too seldom considered.

We developed an open-access, versatile, streamlined, and highly automated pipeline to address these challenges and enable quantitative *tracking* of the *contractile* dynamics of single hiPSC- CMs over time: `ConTraX`. Three interlocking software modules enable: (i) parameter-based localization and selection of single hiPSC-CMs; (ii) automated video acquisition of >200 cells/hour; and (iii) streamlined measurements of the contractile parameters via traction force microscopy. Using `ConTraX`, we analyzed >2,753 hiPSC-CMs over time under orthogonal experimental conditions in terms of culture media and substrate stiffnesses. Using undirected high-dimensional clustering, we dissected the complex diversity of contractile phenotypes in hiPSC-CM populations and revealed converging maturation patterns.

Our modular `ConTraX` pipeline empowers biologists with a potent quantitative analytic tool applicable to the development of cardiac therapies.

---

Cardiomyocytes (CMs) differentiated from human induced pluripotent stem cells (hiPSCs) have emerged as a powerful *in vitro* model for the study of cardiomyopathies, cardiotoxicity and in drug development.^1,2^ However, the most important function of these cardiomyocytes—their ability to contract —remains cumbersome to quantify at high throughput and over time in longitudinal assays.^1^

Current assays of cardiomyocyte contractile function are often mostly qualitative, lack field-wide standards, and generally fail to offer the scalability and throughput necessary for larger-scale studies.^1,2^ For example, high-speed video microscopy of CM contraction is relatively uncomplicated and scalable, but it only provides semi-quantitative measurements of contractile frequency and speed.^2–4^ Other more elaborate approaches, such as multicellular constructs, engineered heart tissue or heart-on-chip systems, quantify the total force produced at the microtissue level, but these data remain difficult to standardize across platforms and studies.^2,5–11^ In contrast, quantitative and standardizable measurements of contractile force at the single-cell level, such as through traction force microscopy (TFM), enable quantifying cell-to-cell variations and population heterogeneity.^3,12–16^ TFM is a conceptually straightforward method that measures the traction stress exerted by a cell on an elastic substrate, often a hydrogel. TFM tracks the hydrogel deformation using fiducial markers, generally embedded fluorescent microspheres.^15,17–22^ The measured deformation is then used to compute the traction stress. Additionally, the stiffness of the hydrogels can be tuned to mimic more physiologic healthy or pathologic tissue mechanics.^19–21^

Despite recent progress,^15,16,23^ TFM data acquisition and downstream analysis remain a demanding, time consuming, and tedious process. The acquisition of video recordings requires several hours per experiment; most of that time spent manually selecting cells of interest; and data processing with currently available TFM computational packages is slow and user-input intensive. These challenges are compounded by the size of the datasets when measuring dynamic contractions in CMs that requires seconds-long, high-framerate videos (5-10 s, 150-300 frames). Consequently, the currently achievable throughput is low, limiting the ability to measure enough single cells and to track changes in the same individual cells over time. These limitations have left the potential of such single-cell resolution measurements largely untapped.

To address these bottlenecks, we developed `ConTraX`, a pipeline that provides an accessible, comprehensive, and streamlined TFM assay workflow. `ConTraX` enables quantitative *tracking* of the *contractile* dynamics of thousands of single hiPSC-CMs over time (**Figure 1**). Using `ConTraX`, we analyzed >3,000 hiPSC-CMs at single-cell resolution to characterize the functional heterogeneity and contractile maturation in hiPSC-CM populations.

**Figure 1:**
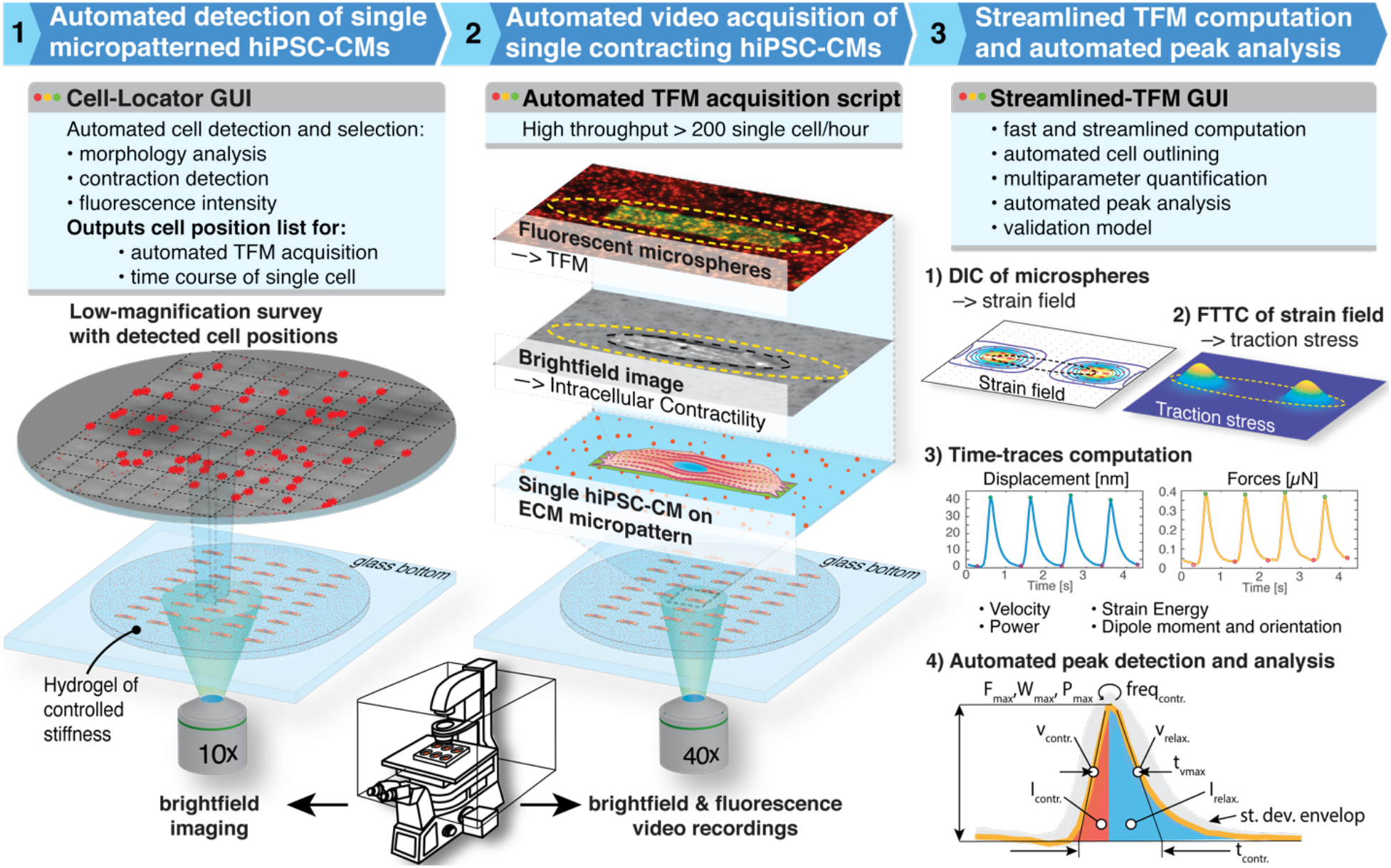
The `ConTraX` workflow is comprised of three main steps and software modules that streamline and automate the workflow, thereby increasing the throughput of the assay. Each module can be used independently or within the workflow. The modules are open-source and available online through our GitHub repository.^24^

## ConTraX workflow

`ConTraX’s` workflow comprises three stand-alone yet interlocking software modules with graphical user interface (GUIs) (**Figure 1**). First, the *Cell-Locator* module (**Figure 1.1**) automates the detection and localization of single cells. Second, the *Automated TFM acquisition* module (**Figure 1.2**) loops through the position list generated by the *Cell-Locator* and automates the recording of high-resolution TFM videos. Third, the *Streamlined TFM* module (**Figure 1.3**) quantifies the contractile phenotype of each recorded cell. The three software modules are open-source and freely available through our GitHub repository.^24^

**Figure 2:**
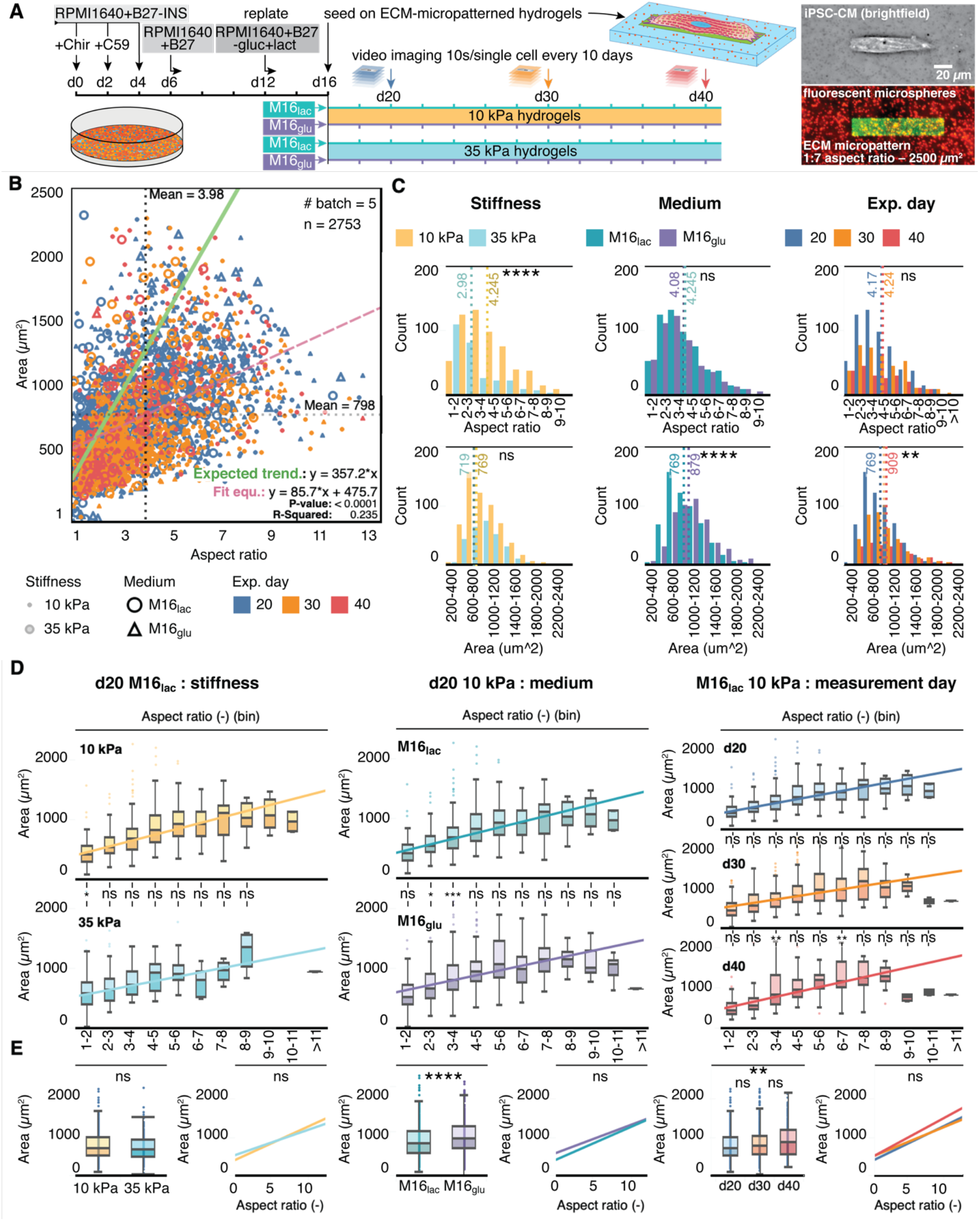
Characterization of cell morphology reveals heterogeneities in populations of hiPSC-CMs. **A) Left:** Schematic of hiPSC-CM culture, differentiation, micropatterning and experimental groups. **Right:** brightfield image of a single micropatterned hiPSC-CM and corresponding fluorescence image; the ECM micropattern (GFP-labelled gelatin + Matrigel) printed on polyacrylamide hydrogel with embedded red-fluorescent. **B)** hiPSC-CMs adopt an elongated morphology when adhered onto the ECM micropatterns (7:1 aspect ratio). Green line: expected trend for the area of a cell filling the whole width of the micropatterned while elongating along the micropattern (Eq. 1). Red dashed line, linear regression; dotted lines: median values for spread area and aspect ratio. **C)** Stiffness, maturation medium, and experimental day influence the distributions of spread area and cellular aspect ratio. **D)** The mean spread areas increases as function of aspect ratio (grouped by range). **E)** Overall mean of spread area depends on stiffness, medium, and experimental day, but the rate of change of spread area as function of aspect ratio (comparison of linear regressions) is the same. In all panels, \**p* < 0.05, \*\**p* < 0.005, \*\*\**p* < 0.001, \*\*\*\**p* < 0.0001. ns, not significant.

**Figure 3:**
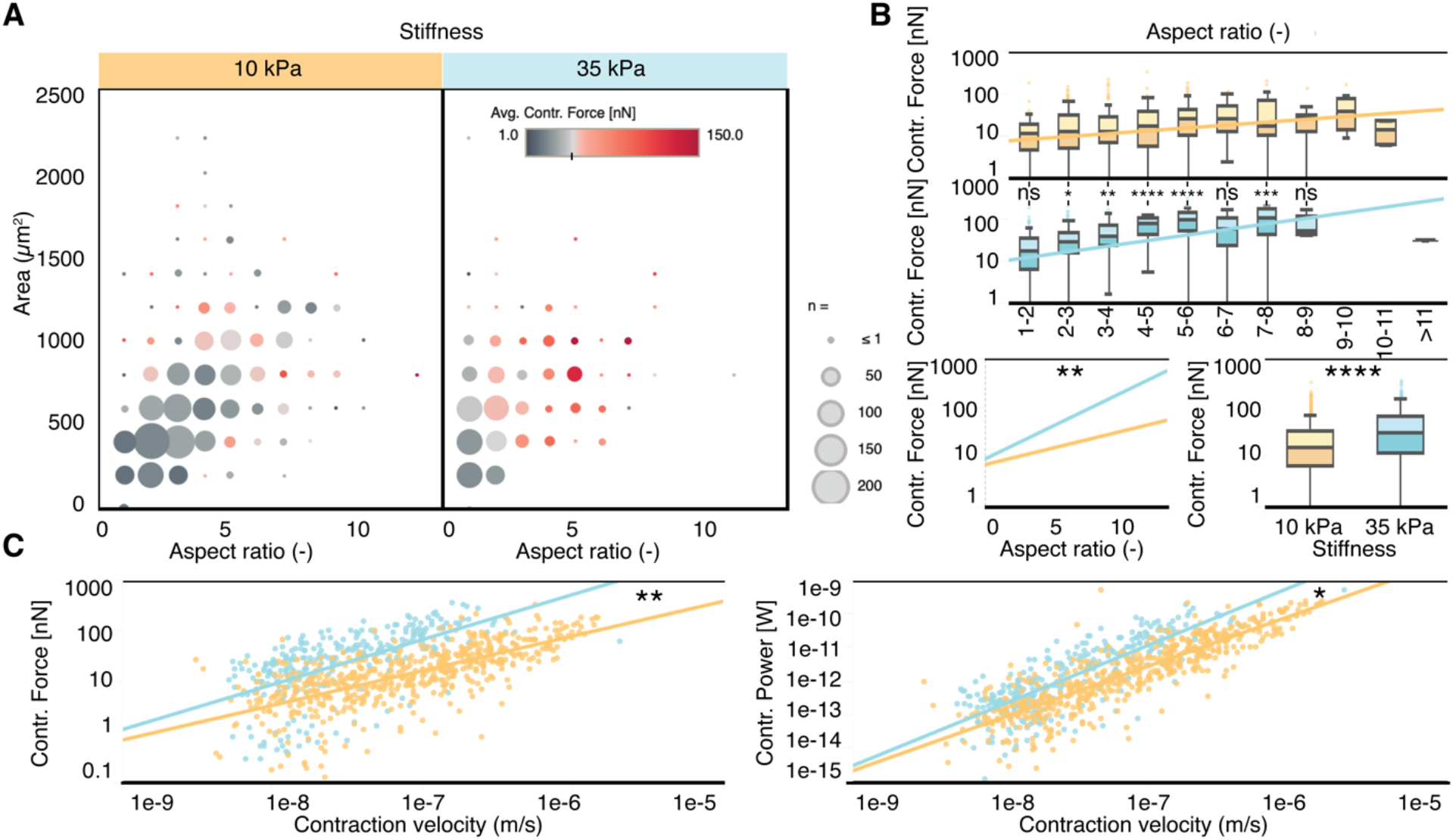
Contraction force, velocity, and power are impacted by substrate stiffness at day 20 in M16lac medium. **A)** Balloon plot of the contractile force as a function of cell aspect ratio and area (grouped by range) shows a generally higher force on 35 kPa substrate and fewer cells of large spread area **B) Top:** Force production increases with aspect ratio until reaching a maximum around aspect ratio 7:1, beyond which force decreases. **Bottom:** Mean force production depends on stiffness and is more dependent on morphology on 35 kPa substrates. **C)** Contraction force and contraction power follow a power law (linear in this log-log plot) with contraction velocity that depends on substrate stiffness. In all panels, \**p* < 0.05, \*\**p* < 0.005, \*\*\**p* < 0.001, \*\*\*\**p* < 0.0001. ns, not significant.

The *Cell-Locator* module uses a low-magnification (10x) survey of the cells on the hydrogel and generates a position list of cells matching user-defined criteria: cell area, elongation, and orientation; and fluorescence intensity and contractility thresholds. This targeted and reproducible selection alleviates user-induced cell selection bias, provides population-wide cell-morphology metrics, and increases the robustness and throughput of the assay. The generated position list also enables tracking the same single cells over time, thereby enabling longitudinal studies.

The *Automated TFM acquisition* module loops over the cell position lists generated by the *Cell-Locator*, moving the microscope stage to the selected cell locations, performing an optional autofocus step, and acquiring video recordings according to user-defined imaging parameters. The module was developed as a script for *Micro-Manager*, a widely used open-source microscopy software, and as a GUI macro for *Zen Blue*, the proprietary Zeiss microscopy software.

The *Streamlined TFM* module, innovating on prior work,^15^ uses Digital Image Correlation (DIC) to track fluorescent fiducial markers (i.e., fluorescent microspheres) and measure the deformation of the hydrogel. Fourier Transform Traction Cytometry (FTTC) is used to compute the traction stress.^12,25^ A custom built algorithm automatically detects contraction peaks and quantifies the contractile parameters: maximum contraction force *F_max_*, work *W_max_*, power *P_max_*, strain energy *E_max_*, frequency *f_contr_*, as well as contraction and relaxation velocities *v_contr_* and *v_rel_*, contraction and relaxation impulse *I_contr_* and *I_rel_*, ^26,27^, contraction durations at maximal velocity, t_vmax_, and overall duration of the contraction, t_contr_. The TFM computation was validated with our validation model and standardization tool,^28^ making the results comparable across labs and studies.

## Results

Substrate stiffness, culture medium and maturation time have been shown to affect cardiomyocyte contractile function by driving differences in cell morphology, intracellular protein structures, and sarcomere assembly and maturity.^29,30^ Therefore, to characterize the contractile phenotype in hiPSC-CMs populations and to demonstrate the power of `ConTraX`, we designed a time-course experiment over three time points (days 20, 30, and 40 post differentiation) across 2×2 orthogonal experimental conditions (two substrate stiffnesses: 10 kPa (healthy myocardium) and 35 kPa (fibrotic myocardium), and two media compositions: M16_lac_: lactate-rich and M16_glu_: glucose-rich) (**Figure 2A**).

Details of the experimental workflow and of the fabrication procedures for the hydrogel substrates are reported in Methods and in the online Detailed Methods. In short, after differentiation^30–33^ from wild-type WTC hiPSC,^34–36^ hiPSC-CMs were seeded on polyacrylamide hydrogel substrates of controlled stiffness, micropatterned by microcontact printing with extracellular matrix (ECM) proteins to constrain the cell morphology.^23,37^ Our micropatterned hiPSC-CMs were left to recover for 5 days, and subsequently imaged and analyzed using `ConTraX`. We measured 3,366 single hiPSC-CMs across five differentiation batches and, after excluding 613 measurements due to cell doublets or unidentifiable contraction peaks (caused by a lack of contraction or low signal-to-noise ratio), our final dataset includes 2,753 single micropatterned hiPSC-CMs. For statistical comparison, our control group is that of cells cultured in M16_lac_ media on substrates with 10 kPa stiffness at day 20 (**Figure 2A**).

### Workflow performance

`ConTraX` workflow enabled unprecedented performance in terms of workflow efficiency and throughput, minimizing the need for intervention and enabling parallelization of experimental and analytical steps. The *Cell-Locator* module could generate a position list for hundreds to thousands of single hiPSC-CMs in only a few minutes. Thereafter, the *Automated TFM acquisition* module enabled a video acquisition throughput of > 200 cells/hour, which is an order of magnitude higher than conventional manual acquisition and significantly reduces the burden on users. Finally, the *Streamlined TFM* module shortened data analysis of this large dataset by > 20x compared to previous tools (on a regular 2015 MacBook Pro laptop).^15^ By further exploiting parallel computing, data analysis was shortened to << 150 s per typical TFM video using a 12-core 2.7 GHz processor workstation.

### Micropatterning controls cell shape

Native adult CMs display well aligned myofibrils and elongated shapes, features that are often not spontaneously developed in largely immature hiPSC-CMs. However, previous reports showed that hiPSC-CMs can adopt more physiological elongated shapes and enhanced contractile maturity when seeded on physiological micropatterned substrates.^18,38^ Therefore, we seeded our hiPSC-CMs on rectangular micropatterns of aspect ratio 7:1 shown to elicit maximal alignment and contractile efficiency.^18,39^ We deliberately chose a micropattern area of ∼2500 μm^2^, larger than the reported average spread area of unpatterned day 30 hiPSC-CMs^22^, to afford us the ability to observe a range of cell morphology while maintaining the necessary spatial constraints to induce cell alignment.

Using the *Cell-Locator* and the *Streamlined TFM* modules, we measured the morphology of our micropatterned hiPSC-CMs as a function of substrate stiffness, culture medium and time (**Figure 2B-D**). Our micropatterned hiPSC-CMs successfully adopted elongated shapes (**Figure 2B**) and were constrained within the micropattern, with a measured mean cell aspect ratio of 3.98 ± 2.02 (median = 3.58) and a mean spread area of 798 ± 383 (std) μm^2^ (median = 733 μm^2^) across all groups at day 20. If all cells filled the whole width of the micropattern as they elongate, the cell spread area would be expected to correlate with aspect ratio following a linear trend defined by: *A* = *L* ∗ *W* = *r* ∗ *W*^2^ (1), where *A* = micropattern area*, L* = length, *r* = length:width aspect ratio, and *W* = 18.9 μm width (for *A* = 2500 μm^2^ and *r* = 7:1) (solid green line in **Figure 2B**). In contrast, our measurements showed a large variance and a smaller slope (red dashed linear regression in **Figure 2B**) compared to (1). Nevertheless, as expected from the dimensional constraints imposed by the micropatterns, cells with an aspect ratio exceeding that of the micropattern (>7:1) had a smaller spread area than the maximum possible 2500 μm^2^ on the micropattern (**Figure 2D**) due to a reduced cell width. Hence, the cells were adequately constrained on the micropatterned and the large variance is mainly a results of cell heterogeneity.

### Cell morphology is affected by stiffness, culture medium and maturation time

The morphology of unpatterned CMs is known to change at increased substrate stiffness *in vitro*, with larger cells developing on stiffness matching that of stiffer fibrotic tissues.^29^ In our micropatterned hiPSC-CMs, a substrate stiffness of 35 kPa resulted in wider and shorter cells compared to control (M16_lac_ on 10 kPa at day 20), with similar spread area but smaller aspect ratio (**Figure 2C** & **Table 1**). CMs also experience changes in their metabolism, switching from glycolysis to mitochondrial oxidative metabolism during early development and reverting to glycolysis under hypertrophic stress.^40–42^ At day 20 on 10 kPa substrates, the glucose-rich M16_glu_ medium resulted in larger cell area than the lactate-rich M16_lac_ medium, with no difference in cell aspect ratio. Over time, however, the M16_lac_ medium led to an increase in the spread area in our M16_lac_ control cells (**Figure 2D** & **E** & **Table 1**), surpassing the M16_glu_ cells by day 30 (**Figure S3**), especially on stiffer substrates. This trend correlates with an increased mitochondrial respiration in M16_lac_ cells compared to M16_glu_ cells at d30, as measured using a Seahorse metabolic assay (see Supplementary Information **Figure S4**).

**Table 1:**
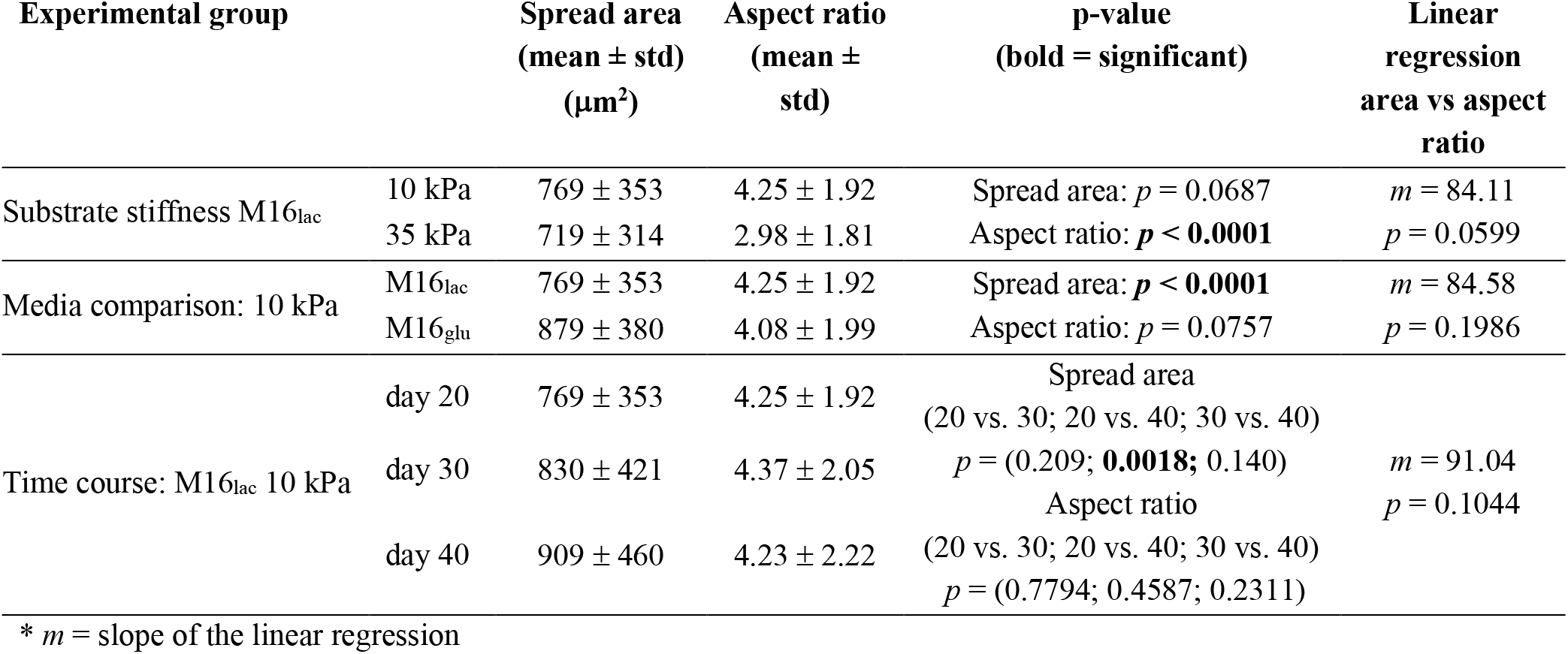
Summary of cell morphology measurements across experimental groups

| Experimental group | | Spread area<br>(mean ± std)<br>($\mu\text{m}^2$) | Aspect ratio<br>(mean ± std) | p-value<br>(bold = significant) | Linear regression<br>area vs aspect<br>ratio |
| --- | --- | --- | --- | --- | --- |
| Substrate stiffness M16<sub>lac</sub> | 10 kPa | 769 ± 353 | 4.25 ± 1.92 | Spread area: $p = 0.0687$ | $m = 84.11$ |
| | 35 kPa | 719 ± 314 | 2.98 ± 1.81 | Aspect ratio: **$p < 0.0001$** | $p = 0.0599$ |
| Media comparison: 10 kPa | M16<sub>lac</sub> | 769 ± 353 | 4.25 ± 1.92 | Spread area: **$p < 0.0001$** | $m = 84.58$ |
| | M16<sub>glu</sub> | 879 ± 380 | 4.08 ± 1.99 | Aspect ratio: $p = 0.0757$ | $p = 0.1986$ |
| Time course: M16<sub>lac</sub> 10 kPa | day 20 | 769 ± 353 | 4.25 ± 1.92 | Spread area<br>(20 vs. 30; 20 vs. 40; 30 vs. 40)<br>$p = (0.209; \mathbf{0.0018}; 0.140)$ | $m = 91.04$ |
| | day 30 | 830 ± 421 | 4.37 ± 2.05 | Aspect ratio<br>(20 vs. 30; 20 vs. 40; 30 vs. 40)<br>$p = (0.7794; 0.4587; 0.2311)$ | $p = 0.1044$ |
| | day 40 | 909 ± 460 | 4.23 ± 2.22 | | |

\* $m$ = slope of the linear regression

### Increased substrate stiffness leads to higher contractile strength

An increased stiffness is also expected to induce CMs to contract more strongly.^29,30^ 35 kPa substrates led to higher contractile force in our control in M16_lac_ at day 20 compared to control 10 kPa substrates (**Figure 3A** & **B** & **Table 2**). 35 kPa substrates also resulted in a higher correlation between contractile force and aspect ratio (or cell area) compared to control 10 kPa substrates (**Figure 3B** and **Figure S1**). Interestingly, both contractile force and contractile power presented the signature of a power-law scaling with contraction velocity (**Figure 3 C** & **Table 2**; see **Figure S2** for comparison of fits), meaning that the contractile force or power scaled as a power of the contractile velocity. An increased 35 kPa substrate also results in a larger slope.

**Table 2.** Summary of contractile force measurements across experimental groups. Control conditions are 10 kPa substrate stiffness, M16lac culture medium, and experimental day 20.

| Experimental group | | Contractile Force<br>(geometric mean &<br>95% conf. int.) (nN)* | p-value<br>(bold =<br>significant) | Slope of linear regression<br>vs Aspect ratio<br>(mean ± SE) |
| --- | --- | --- | --- | --- |
| Substrate stiffness M16<sub>lac</sub> | 10 kPa | 13.88 (12.42 - 15.52) | **$p < 0.0001$** | 3.934 ± 0.776 |
| | 35 kPa | 23.34 (19.54 - 27.93) | | 10.71 ± 1.698 |
| Media comparison: 10 kPa | M16<sub>lac</sub> | 14.16 (12.71 - 15.78) | **$p = 0.0315$** | 3.934 ± 0.776 |
| | M16<sub>glu</sub> | 17.13 (14.93 - 19.68) | | 4.701 ± 1.354 |

\*(lognormal distribution)

|  |  | Power law regression<br>fit p-value, R<sup>2</sup> | Log-log slopes<br>& SE | p-value<br>(bold = significant) |
| --- | --- | --- | --- | --- |
| Substrate stiffness M16<sub>lac</sub> | 10 kPa | Force: **<0.0001**, 0.474 | 0.687 ± 0.0309 | **$p = 0.0029$** |
| | | Power: **<0.0001**, 0.764 | 1.558 ± 0.0365 | |
| | 35 kPa | Force: **<0.0001**, 0.373 | 0.892 ± 0.069 | **$p = 0.0032$** |
| | | Power: **<0.0001**, 0.649 | 1.796 ± 0.0785 | |
| Media comparison 10 kPa | M16<sub>lac</sub> | Force: **<0.0001**, 0.474 | 0.687 ± 0.0309 | $p = 0.0317$ |
| | | Power: **<0.0001**, 0.764 | 1.558 ± 0.0365 | |
| | M16<sub>glu</sub> | Force: **<0.0001**, 0.459 | 0.792 ± 0.0386 | $p = 0.0512$ |
| | | Power: **<0.0001**, 0.698 | 1.677 ± 0.0497 | |

### Glucose-rich culture medium leads to higher contractile force at day 20

We questioned whether the type of metabolic substrate affects the maturation of the contractile function in hiPSC-CMs. At day 20, similar to its effect on cell morphology, the M16_glu_ medium resulted in a higher contractile force than in control M16_lac_ cells (**Figure 4A** & **B** & **Table 2**). However, a change in culture medium did not affect the correlation between contractile force and aspect ratio (**Figure 4B**) (or area, see Supplementary Information **Figure S1**). It also did not meaningfully affect the observed power-law relationship between contractile force or power with the contraction velocity (**Figure 4C**).

**Figure 4:**
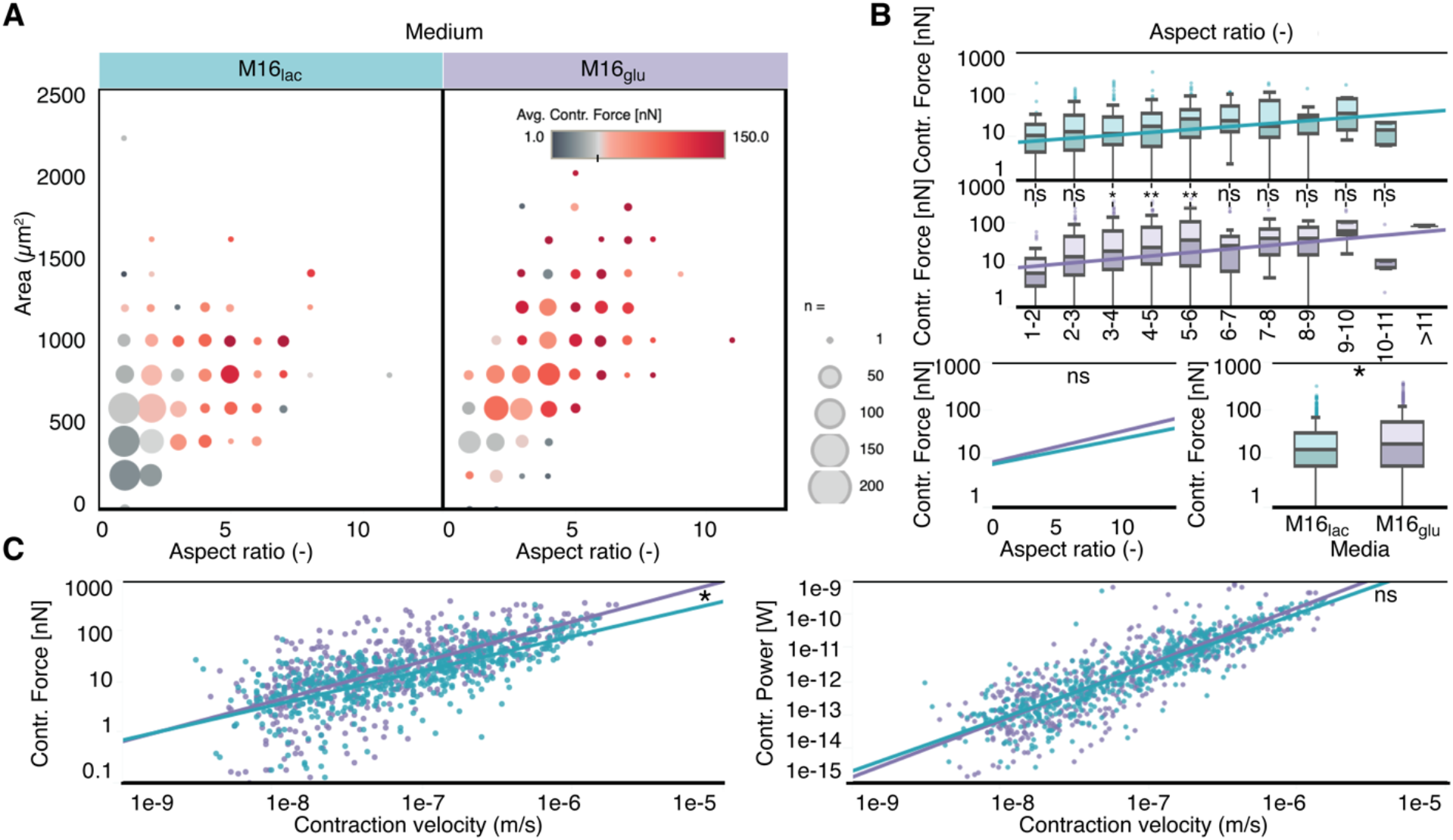
Contraction force, velocity, and power are equally dependent on area and aspect ratio for both medium compositions at day 20 on 10 kPa substrates. **A)** Balloon plot of the contractile force as a function of cell aspect ratio and area (grouped by range) shows more elongated and larger cell in M16glu than in M16lac. **B) Top:** Force is equally dependent on aspect ratio in both media, except in for aspect ratios of 3-6. **Bottom:** Overall mean contraction force is larger in M16glu than in M16lac, but the effect of aspect ratio on force (linear regression) is the same in either media. **C)** Contraction force and power follow a power law (linear in log-log plot) with contraction velocity. The linear regression for the force shows a weak dependence on medium composition. For all panels, \**p* < 0.05. ns, not significant.

### Over time, contractile function is promoted by lactate-rich medium but is attenuated on stiffer substrates

To characterize the maturation of the contractile function over time, we repeatedly measured the same single cells at day 20, 30 and 40 (**Figure 5**). Interestingly, the contractile performance of the M16_glu_ cells declined with time, while that of the M16_lac_ control cells improved (**Figure 5A & B**). The survival of the control M16_lac_ cells was also higher.

**Figure 5:**
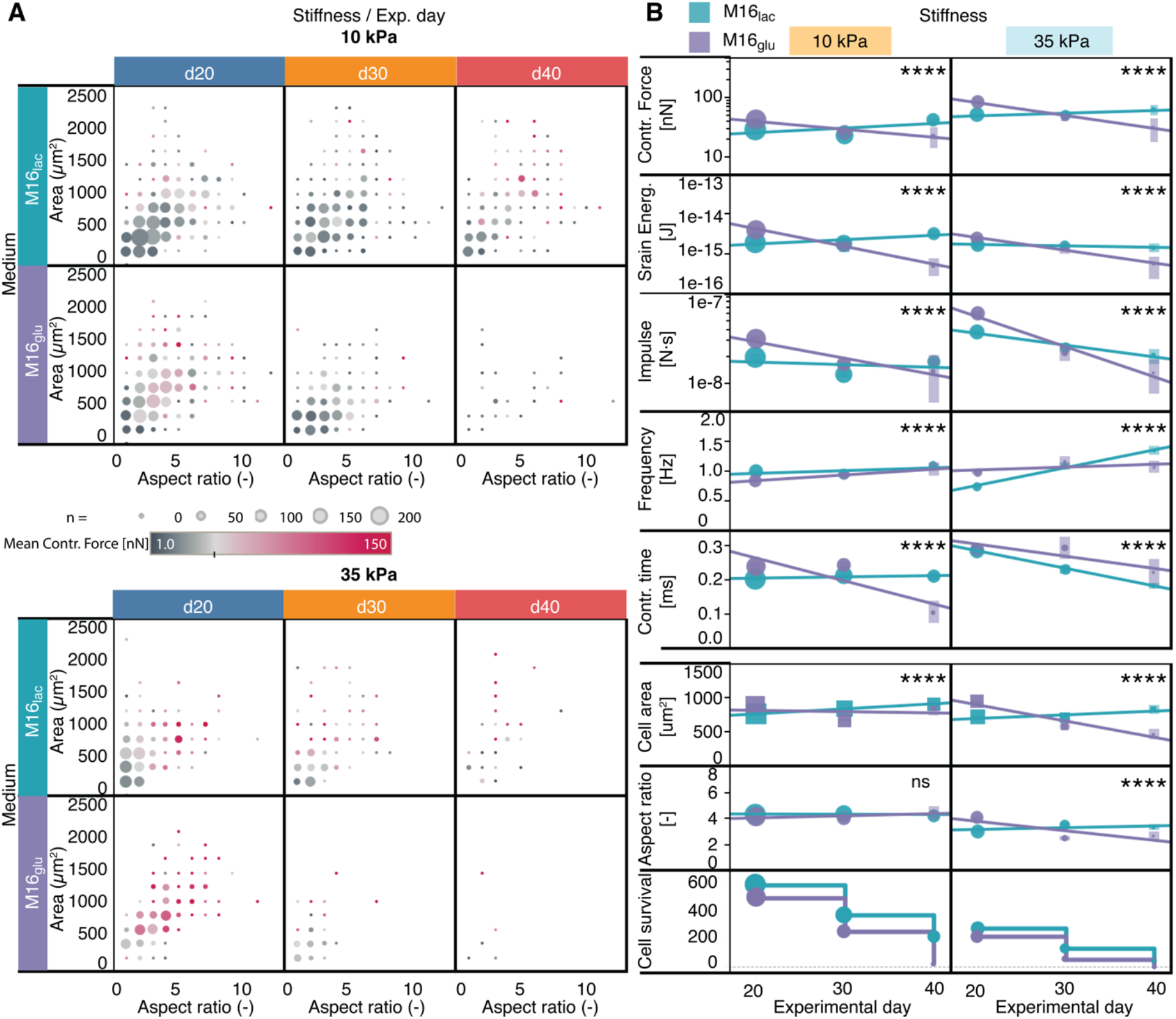
Contractile function in micropatterned hiPSC-CMs increases over time in M16lac versus M16glu medium. **A)** Cells die sooner in M16glu than in M16lac medium and on 35 kPa, while cells in M16lac tend to increase in **B)** `ConTraX` detected changes in various parameters over 20 days, for both medium conditions and substrate stiffnesses. \*\*\*\**p* < 0.0001 for the comparison of the slope of linear regressions; ns, not significant. Vertical bars: standard deviation; lines: linear regressions. See Supplementary information **Figure S3** for more parameters.

Over time, a higher 35 kPa substrate stiffness negatively affected the contractile performance with a plateau (M16_lac_) or decrease (M16_glu_) over time of most contractile and morphology parameters compared to 10 kPa substrates (**Figure 5B**). This attenuation of contractile function on stiffer substrates was also accompanied by faster cell loss or death, which may have resulted from the consistently higher contractile stress observed on these stiffer substrates.

In line with a previously reported progressive loss of rhythmic beating on a rigid substrate,^29^ stiffer 35 kPa substrates led to an increase over time in the mean beating frequency, despite electrical pacing at 1 Hz. Simultaneously, both the total contraction duration and contraction impulse (contractile force integrated over contraction duration) decreased with time (except for the control cells on 10 kPa in M16_lac_ medium).

### Multi-dimensional contractile phenotyping and gating

We leveraged `ConTraX`’s throughput to characterize distinctive multi-parameter contractile profiles in 2,753 single cells using undirected clustering. We used X-shift clustering and identified eight clusters using K-nearest-neighbor density estimation with the following input parameters: average contraction displacement, total contractile force and strain energy, contraction and relaxation velocities and power, contraction duration, beating frequency, cell spread area and aspect ratio (**Figure 6A & C**). (Methods ^43–45^). Clustering most strongly followed from changes in three parameters: contractile force, cell spread area and beating frequency.

**Figure 6:**
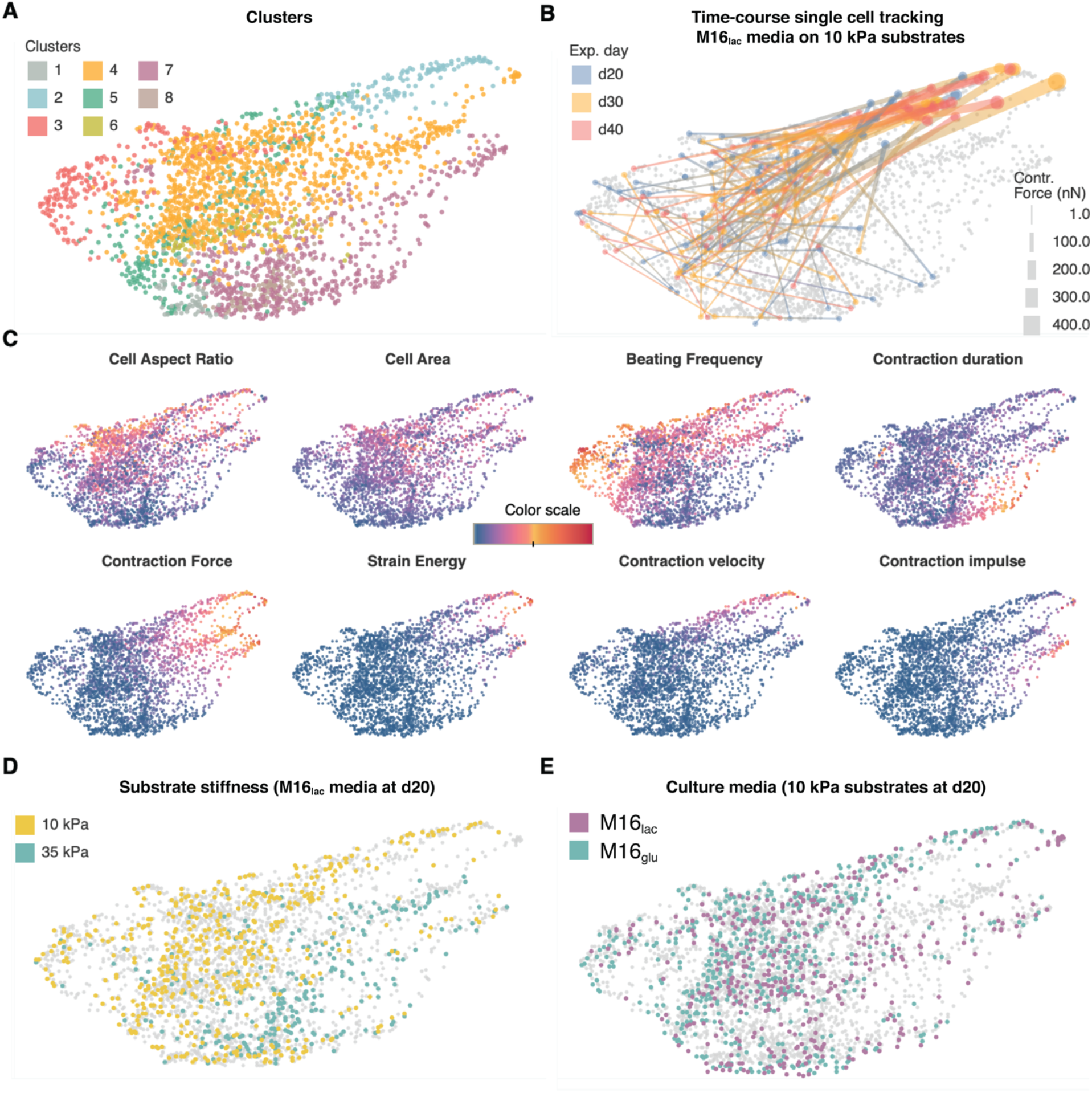
Our large dataset enables cluster analysis, revealing the phenotypic complexity at the single-cell level. **A)** High-dimensional clustering by X-shift K-nearest-neighbor density estimation (Methods) yields eight clusters. **B)** Single hiPSC-CMs in M16**lac** on 10 kPa substrates (control group) over time. Lines connect the different measurement time points for single cells, the color correspond to experimental day, the line width correspond to contractile force. **C)** Distinct distributions of the clustering parameters reveal the characteristics of each clusters. **D)** At day 20, control hiPSC-CMs grown in M16**lac** mainly segregate into clusters 2 and 5 (10 kPa) and into clusters 4 and 8 (35 kPa). **E)** In contrast, control cells at day 20 do not segregate into clusters based on medium composition.

#### Undirected clustering distinguishes distinct contractile phenotypes

Of the resulting clusters (**Figure 6A**), clusters 1, 6 and 8 comprised the smallest cells exerting the least force, while clusters 2, 4, and 7 comprised the largest cells producing the most force (**Figure 6C**). Cluster 5 comprised cells with the least distinct morphology and contractile profile, i.e., a grey zone of the cells most difficult to classify. Cluster 7 comprised the cells with beating frequency < 1 Hz electrical pacing frequency that also displayed a prolonged contraction duration, while cluster 3 comprised the cells with high beating frequency that also displayed a low force production.

Strikingly, clustering identified different contractile phenotypes for the cells cultured on the 10 kPa or 35 kPa substrate stiffnesses, without *a priori* knowledge of the experimental conditions (**Figure 6D**). In line with our earlier observations that stiffness, but not culture medium, impacted contractile function, clustering did not segregate cells cultured in different culture media (**Figure 6D**). Clustering revealed phenotypic differences linked to cell morphology. As expected, the cells with the largest spread areas generally produced the highest force, but, surprisingly, cells with the highest aspect ratios did not necessarily produce a high force. Clustering revealed a phenotypic difference linked to contractile dynamics. Differences in beating frequency and contraction duration resulted in two somewhat opposite phenotypes: one displaying a low force and high beat rate (cluster 3) and another displaying a slow beat rate and prolonged contraction duration (cluster 7). The 1-Hz electrical pacing was most accurately followed by cells with high aspect ratios. Clustering also revealed that the cells producing a particularly high force also displayed a high strain energy, and, interestingly, these cells also displayed a high contraction velocity and were mostly cells cultured on softer 10 kPa substrates. In contrast, other cells contracted with a high force but a slow contraction velocity, and these cells were predominantly large cells cultured on stiffer 35 kPa substrates.

#### Converging phenotype maturation in lactate-rich medium

By exploiting `ConTraX`’s ability to follow single cells over time, we examined the maturation trajectories of the contractile phenotype in individual cells (**Figure 6B**). Notably, the cells in our control group (*i.e.,* cultured in M16_lac_ at 10 kPa substrate stiffness) converged from relatively heterogeneous contractile phenotypes at day 20 (cluster 5) to more homogeneous phenotypes with a larger contractile force (cluster 2). This single-cell resolution analysis confirmed the consistent increase over time in the mean contractile function across individual cells in our control group (**Figure 5B**).

### Subpopulation analysis reveals potential gating strategies

We questioned whether our large dataset and clustering analysis could help identify gating strategies, similar to what is done in Flow Cytometry and Fluorescence-Activated Cell Sorting (FACS) analysis. Here, we divided our dataset in four quadrants based on cell morphologies by defining two gates at the median values of aspect ratio (3.88:1 L:W) and spread area (median = 704 μm^2^) in our control cell group (*i.e.,* cultured in M16_lac_ on 10 kPa substrates at day 20) (**Figure 7A**). Strikingly, a comparison of each quadrant with the clustering revealed that the cells in quadrant 1 almost entirely correspond to those in clusters 2 and 4, the two clusters segregating for phenotypes displaying a large contractile force (**Figure 7B**); and > 80% of the cells in our control cell group had a morphology commensurate with quadrant 1. Interestingly, these cells constituted the bulk of the cells with a converging maturation trajectory (see **Figure 7C**).

**Figure 7:**
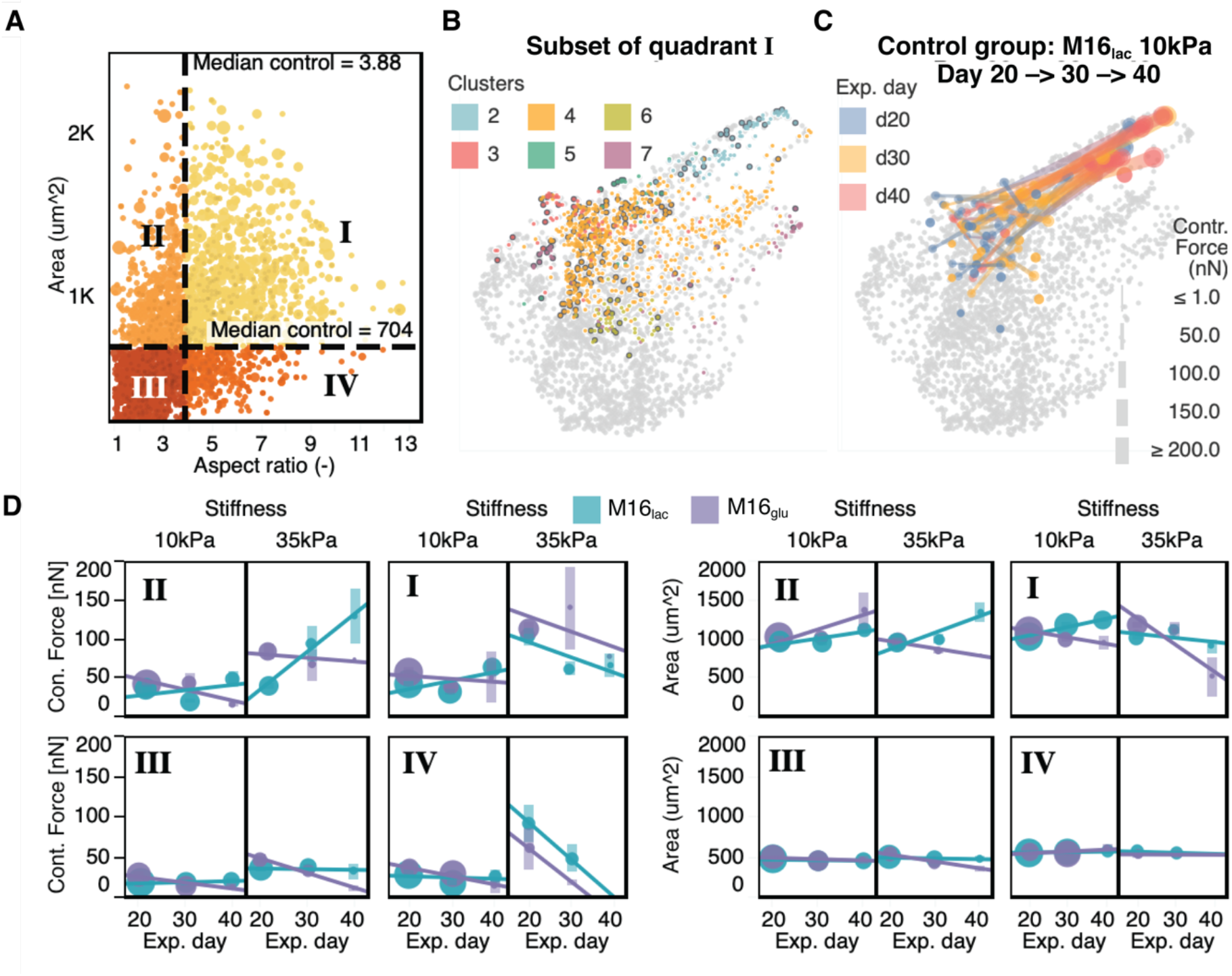
Gating for subpopulations based on cell morphology correlates with difference phenotypic profile. **A)** Similarly to FACS analysis, gates were defined to defined four subpopulations based on cell morphology. The median area and aspect ratio of control cells (M16lac on 10 kPa at day 20) were used as gate values. **B)** Cells in quadrant I correspond closely to cluster 2 and 4 (**Figure 6 A**). Control cells are marked dark borders. Cells in clusters 1, 3, 5, and 7 are almost entirely absent from quadrant I. **C)** Over time, cells in quadrant I progress toward a more homogeneous contractile phenotype. **D)** Despite differences in cell morphology, over time contractile force does not differ across conditions, but the amplitudes and trend differences are amplified in the larger cell (quadrant I and II). Circle size, sample size (n); vertical bars, standard deviation; lines, linear regressions.

We further examined how the trend over time of the mean contractile force and spread area differed in each quadrant (**Figure 7D**). Qualitatively, the trends are generally preserved in each quadrant, with improving contractile function in lactate-rich medium compared to glucose-rich medium (**Figure 5B**). However, as expected, the absolute mean values of the parameters are different depending on the quadrant.

## Discussion

### *ConTraX* workflow offers an order-of-magnitude faster analysis

`ConTraX` enables the acquisition of large datasets with single cell resolution (**Figure 1**). Compared to previous approaches, `ConTraX’s` streamlined workflow and the order-of-magnitude improvement in data processing performance enable assaying hundreds to thousands of single cells within a few days, compared to multiple weeks or months, substantially shortening the time-to-results. Thus, `ConTraX` facilitates the integration of TFM assays in the research workflow for the broader community. Further, `ConTraX`’s interlocking modules afford the ability to track changes in single cells (cardiomyocytes or other cell types as well) over time, a feature that will benefit many longitudinal studies (**Figure 6B**).

### Micropatterning reduces hiPSC-CMs heterogeneity

Micropatterning constrained cell morphology into physiological elongated cardiomyocyte shapes (**Figure 2B**) and contributed to the development of a more mature phenotype, with increasing force with larger aspect ratio (**Figure 3**: Contraction force, velocity, and power are impacted by substrate stiffness at day 20 in M16_lac_ medium. A) Balloon plot of the contractile force as a function of cell aspect ratio and area (grouped by range) shows a generally higher force on 35 kPa substrate and fewer cells of large spread area B) Top: Force production increases with aspect ratio until reaching a maximum around aspect ratio 7:1, beyond which force decreases. Bottom: Mean force production depends on stiffness and is more dependent on morphology on 35 kPa substrates. C) Contraction force and contraction power follow a power law (linear in this log-log plot) with contraction velocity that depends on substrate stiffness. In all panels, \**p* < 0.05, \*\**p* < 0.005, \*\*\**p* < 0.001, \*\*\*\**p* < 0.0001. ns, not significant. and **Figure 4** or area (**Figure S1**).^18^ The micropattern dimensions (1:7 aspect ratio and 2500 um^2^) were intentionally chosen to be larger than that for reported hiPSC-CMs spread areas^18,22^ to enable characterizing the range of cell morphology found in heterogenous hiPSC-CM populations.^46^ As a result, the variance in cell morphology was large. However, the correlation slope between the cell spread area and aspect ratio was not as steep as could be expected from the micropattern dimensions, suggesting that hiPSC-CMs favor elongation over lateral spreading once they begin to elongate (**Figure 2B**). This type of spreading behavior may follow from a more favorable energy balance in the registration of new sarcomere units in series before fusion with neighboring premyofibrils and nascent myofibrils.^47^

### Substrate stiffness and time impacts morphology and contractile phenotype

Stiffer 35 kPa substrates resulted in the development of wider but not smaller hiPSC-CMs over time (**Figure 2C,D** and **E**). This suggests that stiffness and maturation time contribute to promoting cell growth, similar to that reported *in vivo* in ‘compensatory’ hypertrophy.^48^ Stiffer 35 kPa substrates also led to a higher contractile stress (**Figure 3B**), an expected physiological response to increased afterload consistent with previous findings using other *in vitro* models.^17,29,49,50^ Notably, increased stiffness also negatively impacted the contractile performance over time and resulted in faster cell loss or death, revealing a possible link between mechanical stress and loss of contractile function in CMs (**Figure 5B**). Additionally, increased stiffness affected the hiPSC-CMs electrical paceability or beating synchronicity, with a mean beating rate higher than the 1 Hz pacing frequency on 35 kPa substrates. Taken together, these findings suggest an important role of increased tissue stiffness in displacing cardiomyocyte contractile homeostasis, similar to that observed *in vivo* in tissue undergoing fibrosis, as in dilated cardiomyopathy.^26,27^

### A metabolic switch promotes the maturation of contractile phenotype

At day 20, cells cultured in M16_glu_ medium produced more force than control cells (M16_lac_ on 10 kPa at day 20) (**Figure 4**). Over time, however, we observed a dramatic change: the contractile function of cells cultured in M16_glu_ declined and there was a significant increase in cell death, while control cells cultured in lactate-rich M16_lac_ displayed a consistent increase in most contractile parameters (**Figure 5**). This behavior is consistent with the known metabolic switch associated with CM maturation during early development.^40–42^ In accordance mitochondrial respiration increased in control M16_lac_ cells compared to M16_glu_ as seen in Seahorse assays (see Supplementary Information **Figure S4**),

### Undirected clustering identifies relevant phenotypes and maturation trajectories

Our multidimensional clustering analysis (**Figure 6**) provided insight into the diverse contractile phenotypes found in hiPSC-CM populations *in vitro*. `ConTraX`’s ability to measure such phenotypes and maturation trajectories has the potential, together with emerging *in vitro* models of cardiac diseases, to provide insights that can be mapped to *in vivo* observations. For example, *in vivo,* larger cells are characteristic of hypertrophic cardiomyopathy (characterized clinically by hypercontractility), and cells of high aspect ratios are typical of dilated cardiomyopathy (characterized clinically by hypocontractility).^51^ *In vivo,* chamber specificity (atrial versus ventricular) and deficient excitation-contraction coupling have also been linked to differences in contractile dynamics and excitability at the cardiomyocyte level.^39,52^ Further, the well-known force-velocity relationship in the heart and cardiomyocytes links a higher afterload to a lower contraction velocity,^53–58^ This is similar to our observations of a power-law relation between contractile force (or power) and contraction velocity (**Figure 3C & Figure 4C**). Additionally, we demonstrate stiffness-dependent phenotypes: a high force and strain energy is correlated with higher contraction velocity on softer 10 kPa substrates whereas high force with slower contraction velocity is observed in predominantly large cells on stiffer 35 kPa substrates. These findings also reveal a possible allometric power-law relation between contractile velocity and force at the level of the cell. Finally, the larger and slower-contracting cells on 35 kPa substrates displayed a high contraction impulse (the area under the curve of force versus time), a metric that has been suggested as a potential indicator of cardiomyocyte hypertrophy *in vivo* and *in vitro*.^59^

### Unbiased gating potentially allows selection for the cells most able to mature

Our large dataset and clustering analysis revealed how selection gates enable targeting of specific subpopulations. For example, selecting for more elongated and larger cells (quadrant 1) reduces some of the heterogeneity in hiPSC-CMs populations and preselects for the cells that are the most able to further mature over time (**Figure 7**). In `ConTraX`, such morphological criteria can directly be applied as selection gates in the *Cell-Locator* module to preselect subpopulations upstream of TFM data acquisition and analysis. Application of such selection gates is rapid and unbiased in contrast to current approaches that rely on manual and subjective cell selection and further increases the yield in ConTrax and the overall specificity of the assay. Nevertheless, we recommend careful definition and application of morphological selection gates, as these choices can lead to inadvertent exclusion of certain phenotypes of interest, such as high contractile frequency (cluster 3) or hypercontractile phenotypes (cluster 7) of particular importance for certain longitudinal drug or disease studies.

## Outlook

In summary, `ConTraX` combines an innovative approach with three interlocking software modules for multiparameter quantification of the contractile function of single hiPSC-CMs over time. Our analysis demonstrates the power of `ConTraX` in tracking changes in contractile phenotypes with single cell resolution over time. `ConTraX` is a powerful new tool that will benefit longitudinal studies of cardiac contractile deficiencies and contribute to the development of cardiac therapies and personalized medicines.

## Methods

### Extended Methods appear in the Online Data Supplement and data are available via Open Science Framework (OSF) (https://osf.io/785vp/).^60^

### Cell culture and assays

hiPSC-CMs were differentiated in five differentiation batches prepared from different passage numbers of WTC-iPSCs line (one differentiation batch from p37, one from p41, two from p46, one from p47).^34,35^ hiPSCs were differentiated into beating CMs according to established protocols (online Detailed Methods and **Figure 2 A**).^61^ At day 12, the hiPSC-CMs were subjected to 4 days of glucose starvation to purify against fibroblasts, then were replated onto freshly prepared Matrigel-coated tissue culture plates. At day 16±1 on average, half of the cultured wells were either switched to RPMI-1640 + B27 + D-glucose medium or maintained with RPMI-1640 + B27 + DL-lactate medium.^40^ These media are called M16_glu_ (glucose-rich) and M16_lac_ (lactate-rich), respectively, in the rest of this report. On day 17, the hiPSC-CMs were replated onto micropatterned hydrogel devices for TFM. Cells (∼20 x 10^5^ cells/cm^2^) were replated using the medium in which they were last cultured plus 5 μM ROCK inhibitor (#Y-27632, StemCell Technologies) and 5% Gibco™ KnockOut™ Serum Replacement (KSR, #10828010, ThermoFischer Scientific). Medium was replaced, without ROCK inhibitor and KSR, after 24 h and thereafter every 2 days. Cells were allowed to recover for at least 48 h after seeding before measurements.

### Time course of contractile force maturation of single hiPSC-CMs

To evaluate the power of `ConTraX`, we designed a 2-dimensional time course experiment that aimed to reveal the effects of stiffness and of culture-medium composition on the contractile maturation of micropatterned hiPSC-CMs over 20 days (**Figure 2 A**). We employed two substrate stiffnesses, 10 kPa and 35 kPa, which correspond to healthy and fibrotic tissue, respectively.^29^ For consistency, all hydrogel devices were prepared in batch two days before replating the hiPSC-CMs, using the same stock solution of hydrogel precursors and reagents to minimize experimental variability. Micropatterns of ECM protein with a 2500 μm^2^ rectangular area and a 7:1 aspect ratio were defined via microcontact printing and transferred onto the hydrogel via copolymerization, spatially confining single hiPSC-CMs into a physiologically relevant, elongated shape.^62^

On days 20, 30, and 40 after initial differentiation (day 0), TFM videos of single cells were acquired using `ConTraX`. For each cell batch, 10x-magnification surveys and 40X-magnification traction force videos were acquired for all conditions on the same day. Cell were electrically paced at 1 Hz using a bipolar pulse of 20 V and single-well carbon electrodes during video acquisition. A 5% CO_2_, 37 °C, full stage enclosure incubation chamber was used on the microscope setup to ensure optimal environmental conditions.

At d20, we used `ConTraX`’s *Cell-Locator* module to locate single micropatterned hiPSC-CMs. On subsequent imaging days, we reused the position list from this initial device survey to automatically locate the same single cells and image them again. To use the *Cell-Locator* module, we acquired tiled image surveys of our micropatterned hiPSC-CMs on the TFM substrates using bright field microscopy (transmitted light) and a 10X, 0.3 NA air objective, using the built-in *Acquire Multiple Regions* plugin of *Micro-Manager* software v1.4 with autofocusing (**Figure 1**, middle). The *Stage Position List* of the absolute stage XYZ coordinate for each tile was saved as a position file (.pos) file. Tiled images (.tif or .czi) and the corresponding position list were loaded into our *Cell-Locator GUI*. The software identified single cells were automatically based on user-defined selection criteria. For this study, we defined broad search criteria (cellular aspect ratio of 1.5:1 to 12:1 (length:width) and an area of 300-3500 μm^2^) to assay the heterogeneity of the hiPSC-CMs on our micropatterned substrate. The *Cell-Locator* software module outputs a position list of all the cells meeting these criteria. Details of the workflow in the *Cell-Locator* module are found in the online Detailed Methods.

Next, we used `ConTraX`’s *Automated TFM acquisition* module to automatically acquire TFM videos of these cells. We first calibrated the positions for any x-y-z offset resulting from differences between objectives or stage drift. We recorded 7-s videos using a 40X, 0.6 NA air objective, with a 3-ms exposure for the bright field channel and a 20-ms exposure for the fluorescence channel, using stream acquisition (the maximum camera speed for a given exposure). The dose of fluorescence illumination was minimized to limit phototoxicity. We applied a predefined crop factor of 600 x 300 pixels (no pixel binning). These parameters yielded an acquisition frame rate of 30 frames/s on our setup, which is above the minimum necessary rate (>15 frames/s) to avoid undersampling and adequately quantify of the contractile dynamics of CMs. At each cell position, software autofocusing was performed using the ‘OughtaFocus’ as *Autofocus properties* in *Micro-Manager*, or with the software autofocus in *Zen* (Zeiss) Settings were optimized for our experimental conditions (search range 40 μm; tolerance 1 μm; crop factor 0.5; exposure 1 ms; default lower and upper FFT cutoffs of 2.5% and 14%, respectively; maximizing for Edges detection).

Finally, we analyzed TFM videos using `ConTraX`’s *Streamlined-TFM* module. Our *Auto-*draw function enabled automated detection of cell outlines. We used the corresponding video parameters (frames per second = 30, pixel-to-micron conversion = 0.275) and applied our built-in cropping and 2-by-2 binning functions to reduce file size using our default 200×85 μm cropping window, leaving enough space around each cell to capture substrate deformation far from the cell itself. These is important for the accuracy of TFM analysis. We used the following displacement parameters: subset radius = 15 px, spacing coefficient = 5 px, cutoff = 10^6^, maximum iteration = 20. For computation of TFM via FTTC, the regularization parameter was automatically computed by the software using L-curve identification.^63^ Contraction peaks are detected automatically using our built-in peak detection and averaging algorithm. Details of the *Streamlined-TFM* module workflow appear in the Detailed Methods online.

### Statistics

Data analysis and statistical analyses were performed with *Tableau Desktop 2019.1* (Tableau Software Inc.) and *Prism 8* (GraphPad Software LLC). Data were tested for normality versus lognormality using the Anderson-Darling test, the D’Agostino-Pearson omnibus normality test, the Shapiro-Wilk normality test, and the Kolmogorov-Smirnov normality test with Dallal-Wilkinson-Lillie for P value; the appropriate log transformation was applied to the data when lognormality was statistically motivated (p < 0.05). Population means were compared using two-tailed parametric t-tests, one-way analysis of variance (ANOVA) for unpaired measurements plus Tukey’s correction for multiple comparisons, or two-way ANOVA with correction for multiple comparisons using Sidak hypothesis testing. *p-*values < 0.05 were considered significant. Least-square regressions were performed on the data without constraints and slope differences between treatments were compared using a sum-of-square F test with significance at *p* < 0.05.

Single-cell clustering was performed in the *Vortex clustering environment*, a freely accessible software that automatically defines high-dimensional clusters within single-cell datasets.^43–45^ We used *X-shift* gradient assignment by Euclidean distance and weighted k-nearest-neighbor density estimation with the following data fields as clustering parameters: cell area, cell aspect ratio, average microsphere displacement, total force, contraction and relaxation velocities, strain energy, contraction and relaxation powers, contraction and relaxation impulses, and contraction frequency and duration. The optimal number of clusters was determined via the built-in elbow detection, which was found at the free parameter value K = 20.

## List of Abbreviations

hiPSC: Human-induced pluripotent stem cell
CM: Cardiomyocyte
TFM: Traction force microscopy
PIV: Particle image velocimetry
DIC: Digital image correlation
FTTC: Fourier transform traction cytometry
GUI: Graphical User Interface

## Author contributions

G.P., H.L., A.S.V.R., A.C.Y.C., H.M.B., and B.L.P. conceived and designed experiments. G.P., H.L., and F.B. developed and designed the computational algorithms and software. G.P. and H.L. carried out the experiments. G.P., H.L., and K.K. carried out the computation. G.P., A.S.V.R., E.C., C.B., and R.W. helped with cell culture and with testing of algorithms. G.P., A.S.V.R., E.C., and B.L.P. analyzed data. All authors contributed scientific insights and to the writing of the manuscript.

## Acknowledgments

We thank all members of the Pruitt laboratory at Stanford University and UC Santa Barbara and members of the Blau laboratory at Stanford University for helpful discussions and support. Wild-type hiPSCs (WTC cell line) were obtained as a generous gift from collaborators, and from the Allen Cell Collection.^34–36^

## 1. Detailed Methods

### Fabrication of micropatterned hydrogel substrates

To manufacture our microphysiological assay substrates, we adapted previously published procedures.^18^ In short, polydimethylsiloxane (PDMS, Sylgard^®^ 184 Silicon Elastomer Kit, Dow Corning Corporation) was molded onto a microfabricated SU8-silicon mold by mixing the PDMS kit components at a 1:10 ratio of curing agent:prepolymer using a Thinky mixer, desiccating for 60 min using house vacuum, and curing at 60 °C for at least 60 min to obtain 1-cm^2^ stamps able to print >4000 patterns of 132.3 μm x 18.9 μm ≅ 2500 μm^2^ and aspect ratio 7:1. Stamps (prechilled at 4 °C) were incubated at 4 °C overnight with Matrigel extracellular matrix (ECM) proteins diluted 1:10 in L-15 medium (Leibovitz), rinsed once with L15 medium, and dried under nitrogen flow. ECM protein micropatterns were microcontact-printed onto cleaned (O_2_ plasma, 80 W, 60 s) 15-mm coverslips by gently placing the Matrigel-coated stamp face down onto the glass, applying a constant pressure with a 50-g weight for 3 min, leaving the device without weight for another 2 min, then separating the coverslip from the stamp using forceps.

Polyacrylamide gel precursor solutions were prepared by mixing acrylamide (CAS # 79-06-1, Sigma) (10% w/v), *N*,*N*′-methylenebisacrylamide (CAS # 110-26-9, Sigma) (0.1% or 0.3% w/v for 10 kPa and 35 kPa, respectively), 3 mM HEPES (Life Technologies), 0.2 µm yellow-green or red fluorescent FluoSpheres™ Carboxylate-Modified Microspheres (#F8813, Invitrogen) (2.16% w/v to yield a final concentration of ∼6 × 10^9^ microbeads/mL), and Milli-Q water and desiccating the solution for 30-60 min in vacuum. While the solution was desiccating, glass-bottom 6-well plates (MatTek Corp.) were silanized with 0.3% 3-(trimethoxysilyl)propyl methacrylate (CAS # 2530-85-0, Sigma) in 200-proof ethanol, adjusted to pH ∼3.5 with 5% acetic acid glacial for 5 min, washed twice with 200-proof ethanol, and dried with N_2_ gas. Polyacrylamide hydrogels were formed by adding ammonium persulfate (Sigma) (0.1% w/v) and *N*,*N*,*N*′,*N*′-tetramethylethylenediamine (Sigma) (0.1% v/v) to the precursor solution, rapidly transferring 35 μL onto a silane-treated glass-bottom well, and placing a microcontact-printed coverslip face down onto the polymerizing gel precursor. The use of 6-well (or larger) plates enables optimal repositioning of the plate onto the microscope stage for acquisition of the same single cell during time-course experiments. In order to obtain reproducible hydrogel substrates, PDMS or Kapton^®^ spacers of the desired thickness were placed between the bottom glass and the microcontact-printed coverslip to obtain a controlled hydrogel thickness of at least 100 μm. At smaller thicknesses, the human induced pluripotent stem cell-cardiomyocytes (hiPSC-CMs) would feel the rigid glass bottom, yielding inaccurate measurements of traction force. After 30 min for polymerization at room temperature, the wells were flooded with phosphate-buffered saline (PBS; pH 7.2, Gibco) plus 1:100 penicillin-streptomycin (10,000 U/mL) (#15140122, Thermo Fisher Scientific) and incubated overnight at 37 °C. Next, the top coverslip was removed to reveal the ECM micropattern transferred into the top surface of the hydrogel. The PBS was aspirated, and hiPSC-CMs were seeded onto the micropatterned hydrogel, as described in the Method section.

### Culture, passaging, and differentiation of hiPSC-CMs

Here we used hiPSC-CMs as an *in vitro* model of human cardiac biology. WTC hiPSCs were obtained as a generous gift from collaborators,^34,35^ and from the Allen Institute cell collection, purchased through the Coriell Institute Biobank. WTC hiPSCs were cultured in Gibco™ Essential 8™ Medium (#A1517001, Thermo Fisher Scientific) supplemented with E8 supplement with daily medium change with Falcon multi-well (6 or 12) culture plates (#353046, #353043, Corning). These plates were previously incubated for at least 1 h with Corning™ Matrigel™ GFR Membrane Matrix (#CB-40230, Fischer Scientific) diluted 1:100 in Leibovitz’s L-15 Medium (#11415064, Thermo Fisher Scientific) at 4 °C. hiPSCs were passaged at 1:30 dilution every 3-5 days or before reaching 90% confluency using 0.5 mM ethylenediaminetetraacetic acid (EDTA) in PBS pH 7.2 (#10010023, Thermo Fisher Scientific) or Accutase solution (#A6964, Sigma-Aldrich). Before passaging hiPSCs, the Matrigel incubation solution was aspirated from a new culture plate and cells were passaged into E8 medium supplemented with 5 μM Rock inhibitor (#Y-27632, StemCell Technologies).

hiPSCs were differentiated into beating CMs according to established protocols.^61^ When hiPSCs reached ∼60-80% confluency (day 0), E8 culture medium was replaced with RPMI-1640 culture medium containing L-glutamine and glucose (#11875093, Thermo Fisher Scientific) supplemented with B-27™ Supplement minus insulin (#A1895601, Thermo Fisher Scientific) and 6 μM GSK-3α/β inhibitor CHIR-99021 (Chir, #S2924, SelleckChem). Forty-eight hours after the start of differentiation, the culture medium was replaced with fresh medium in which Chir was replaced with 2 μM Wnt-C59 (C59, #S7037, SelleckChem), a PORCN inhibitor for Wnt3A-mediated activation. On day 4, the medium was exchanged again with fresh RPMI-1640 without any inhibitor or activator. On day 6, the medium was exchanged again with fresh RPMI-1640 with L-glutamine and glucose (#11875093, Thermo Fisher Scientific) supplemented with B-27™ Supplement (50X) and without serum (#17504001, Thermo Fischer Scientific); the medium was exchanged again on day 8 with the same medium.

On days 10-12, robust beating of the cell monolayer was typically obtained (data not shown) and differentiated hiPSC-CMs were passaged at 1:4 or 1:6 into new 12-well plates freshly coated with Matrigel. Accutase (0.5 ml in 12-well plates, 1 ml in 6-well plates) was used to detach cells, then quenched with RPMI 1640 culture medium with L-glutamine and no glucose (#11879020, Thermo Fisher Scientific) supplemented with B-27™ Supplement (50X) without serum (#17504001, Thermo Fischer Scientific), 4 mM sodium DL-lactate solution (#L7900, Lot#LBR6294V, Sigma-Aldrich), and 5% Knockout Serum (#10828010, Thermo Scientific). This solution was spun for 3 min at 300 x *g* at room temperature, and cells were resuspended in 1mL of the same medium supplemented with 5 μM Rock inhibitor (#Y-27632, StemCell Technologies). Cells were replated onto a new plate, and the medium was changed after 2 days to remove the Rock inhibitor. Spontaneous beating reoccurred soon after replating (data not shown).

### Live-cell microscopy

We used a Leica DMi6000b epifluorescence microscope equipped with a motorized x-y stage, a motorized focus drive, motorized objectives, motorized filter cubes, a motorized condenser turret, a high-speed/sensitivity Photometrics PRIME 95B sCMOS camera, and a live-cell incubator enclosure to maintain optimal cell viability at 37 °C and 5% CO_2_ supplied from a premixed air-CO_2_ gas bottle (Praxair) during imaging. The free and open-source *Micro-Manager* (version 1.4.23 64bit) software was used to control microscope acquisition. The multi-well plate format allowed for culturing cells over long periods in well-controlled conventional cell-culture incubators, while reproducibly repositioning assay substrates on the imaging platform for time-course measurements.

During acquisition, the hiPSC-CMs were subjected to electrical pacing using a single-channel Myopacer Cell Stimulator (IonOptix) and a custom single-well large-area carbon electrode (IonOptix), with a programmed bipolar pulse of 20 V at a frequency of 1 Hz. For video imaging, we used an acquisition cycle of either ∼23 s/cell (autofocus, 7 s brightfield imaging, 7 s fluorescence imaging, and microscope control overhead), ∼26 s/cell (autofocus, a brightfield snapshot, 7 s fluorescence imaging, and microscope control overhead—only a single bright field image is strictly required for cell outlining, not an entire movie).

### Seahorse assay

hiPSC-CMs were seeded at 30,000 per well in Matrigel-coated Seahorse plates in M16_lac_ or M16_glu_ medium five days prior to assay. The oxygen consumption rate responses of cells were measured with the Seahorse Bioscience XF96 flux Analyzer following instruction in the XF cell Mito Stress Test Kit User Guider. All measurements of oxygen consumption rate (OCR) were acquired at 5-min intervals with 1-min mixing between intervals. Three baseline measurements were acquired followed by injection of oligomycin to a final concentration of 2.5 μM. After three measurements in the presence of oligomycin, FCCP was injected to a final concentration of 1 μM and three measurements were recorded. Last, rotenone and antimycin A were injected to a final concentration of 2 μM each, followed by three measurements. CMs were subjected to the same substrate conditions as for culture (M16_lac_ or M16_glu_ medium) but prepared with Phenol red-free unbuffered RPMI-1640 (Agilent) as basal medium instead of RPMI-1640 (Thermo Fisher Scientific). OCR was normalized to live cell count using a PrestoBlue dye in accordance with the manufacturer’s protocol (10 min incubation at 37 °C) and quantified on a TECAN Pro1000 plate reader (Stanford High-Throughput Bioscience Center) at 560 nm excitation and 590 nm emission.

### Software: detailed description and workflow

#### Cell Locator

##### Description

The `ConTraX` *Cell-Locator* module is a MATLAB-based graphic user interface (GUI) that enables the automated selection and localization of micropatterned hiPSC-CMs. We developed it to increase the throughput of video acquisition by removing the bottleneck created by the manual localization and identification of single contracting hiPSC-CMs (**Figure 1.1**). Using a sequence of image-processing steps including thresholding, edge detection, masking, and filtering, thousands of relevant single hiPSC-CMs are automatically identified and located within tens of seconds. User-defined morphological search criteria filter subpopulations and exclude alien objects; contraction detection and fluorescence expression level can be used as additional filtering criteria. Automation using such criteria increases the yield of TFM acquisition by only selecting contracting cells or identifying a cell subtype based on the level of expression of a specific protein using a live-cell fluorescent reporter or dye. The absolute positions of the cells on the microscope stage is calculated from their respective locations within a tile image and the morphology of each cell is analyzed in terms of area, width, length, aspect ratio, and main elongation axis orientation.

As such, this software leads to rapid and unbiased selection and morphological analysis of single cells, and—critically—enables the subsequent automated acquisition of TFM video recordings. The software’s output is of broad applicability to the analysis of cell morphology in any given cell population and single-cell detection, and not solely limited to the analysis of contracting hiPSC-CMs via TFM.

##### Input – Output

The *Cell-Locator* module takes as input images of file types *.tiff* or *.czi*, forming the tiles of the survey. If contraction detection or fluorescence intensity is desired for detection or analysis, then time stacks or multichannel stacks can be provided as input. For *.tiff* images, the associated stage position must be provided, which can be obtained from the microscopy software used for survey acquisition. Any desired magnification and illumination technique (*e.g.,* bright field, phase, or fluorescence microscopy) can be used, and the corresponding pixel-to-micron conversion must be provided.

The *Cell-Locator* module outputs: 1) a position list for subsequent automated acquisition of TFM videos at higher magnification; and 2) a results file with single cell-level morphological information and population statistics. The x-y-z positions of the cells are returned as a position list *.pos* file for *Micro-Manager* (The Open Source Microscopy Software: https://micro-manager.org) or as a stage mark list *.czstm* file for *Zen Blue 2.6* (Carl Zeiss Microscopy: https://www.zeiss.com/microscopy/us/products/microscope-software/zen.html); results are exported to a *.csv* spreadsheet.

##### Workflow

After providing the input data, the user is requested to define search criteria for the ranges of area, aspect ratio, and orientation in which cells are to be identified and selected. Before automatic processing of the entire stack of images of the survey, the search criteria can be quickly tested and refined using a single image. The software then automatically detects thousands of single cells matching the search criteria. If desired, the detected cells can be reviewed and excluded manually, and additional cells can be manually added. Identified cells are marked in each tile image and a zoomed-in image of each cell with its detected outline is displayed on the side of the screen with the corresponding morphological parameter for easy and rapid review.

If contraction detection is desired, then time-stack images can be provided as input and a built-in algorithm detects cell contraction for further filtering. To achieve this detection, the image survey can be performed using a short time lapse of >2 frames at a framerate sufficient to slightly oversample a contraction cycle. For example, 4 frames at 166-ms intervals maximizes the chance of acquiring at least a part of the contracted or relaxed state of a cell beating at 1 Hz. Our time-efficient algorithm calculates the z-project standard deviation for each pixel, as a measure of the local variation over time. It then calculates the average value of this standard deviation within a cell area and determine whether the local variation over time within the cell area is above a user-chosen threshold compared to the rest of the frame (for example how the average time standard deviation of a cell compares to the average time standard deviation in the image). A slider adjusts the threshold value and the software dynamically displays the threshold mask over the image, enabling easy identification of an optimal threshold for a given image.

If fluorescence-based selection is desired, then multichannel stack images can be provided as input and a built-in algorithm filters cells based on fluorescence intensity. In this case, the image survey should be performed with the appropriate fluorescent channel in addition to the transmitted light imaging. The average intensity of fluorescence is calculated within each cell area and a slider enables the user to adjust the detection threshold (similar to contraction detection), thereby enabling selection of cells based on fluorescence intensity.

#### Automated TFM Acquisition module

##### Description

The *Automated TFM Acquisition* script automates the acquisition of TFM videos of single contracting hiPSC-CMs. We developed it to address the throughput bottleneck created by the manual acquisition of live videos of previously identified single contracting hiPSC-CMs. *Automated TFM Acquisition* contributes to increasing the throughput of the workflow and reducing the burden on user/s by automating the acquisition of video recording using the cell position list from the *Cell-Locator* module (**Figure 1.1**). Two versions were developed, one each for *Micro-Manager* and for *Zen Blue 2.6*, because neither software enables a direct acquisition sequence in hierarchical order: *Position (with autofocus at each position) à Channel à Time (video streaming)*. The scripts were written in the *Beanshell* scripting language in *Micro-manager*, and in *Python* in *Zen*. Both versions are fully editable and can be rapidly adapted to fit specific user requirements.

##### Input–Output

*Automated TFM Acquisition* takes the cell positions list from the *Cell-Locator* module as input; these positions are loaded as *Stage Position List* in *Micro-Manager* and as *Stage Mark* in *Zen*. The script automates acquisition by using the acquisition parameters (such as exposure time) that the user defines in the microscopy software during experimental setup. During execution, the script loops through the positions list to acquire videos of each hiPSC-CM, first using transmitted light and then using the fluorescence channel corresponding to the fluorescent microspheres (**Figure 1.2**).

The software outputs videos saved in 8-bit *\*.tiff* format directly in order to minimize data volume or as *.czi* files in *Zen*. Each file name retains the identity of each cell for easy pairing of single-cell data in time-course experiments.

##### Workflow

After loading the position list from the *Cell-Locator* module and setting up the experimental parameters in the microscopy software, the user executes the *Automated TFM Acquisition* script. If the culture plate has already been returned to the incubator, it can be loaded again onto the microscope stage. In this case, the cell position list may be off by a fixed offset in x, y, or z; the user should check for an offset, and correct it, within the microscopy software. During automated acquisition of a large number of cells, and due to possible unevenness of the hydrogel surface, we recommend that the user includes a software autofocus step at each cell position (this is an option in *Automated TFM Acquisition*) to ensure adequate focus. In addition, a focus offset can be programmed to adjust for differences in focal plane between bright field and fluorescence microscopy. Accurate TFM measurements require a good focus position at the top surface of the hydrogel (in direct contact with the cells) when measuring the displacement of fluorescent microspheres.

An additional version of this script yields semi-automated acquisition, making it possible for the user to skip positions or to navigate along the position list manually, with rapid keyboard strokes. This semi-automation helps reduce the overall experimental time at later acquisition timepoints during a time course, as many cells may have detached or died by these later timepoints.

#### Streamlined TFM

##### Description

*Streamlined TFM* is a MATLAB-based GUI for analysis of TFM video data. It addresses the throughput bottleneck created by the computationally- and user-intensive process involved in the processing of TFM data. The *Streamlined TFM* contributes to increasing the throughput of the workflow by reducing the need for user input during data computation and analysis and by speeding up the computation by exploiting parallel processing and efficient data handling (**Figure 1.3**). The *Streamlined TFM* was developed building upon the work of Ribeiro et al.^15^ as the base code architecture and TFM algorithm implementation.

*Streamlined TFM* offers numerous features in addition to significant streamlining and more efficient computing. Of notable importance are two new algorithms for automated cell outlining and automatic contraction peak analysis. Both these algorithms reduce the amount of user input (mouse clicks) by at least an order of magnitude, save a considerable amount of time in terms of data pre/postprocessing, and, importantly, reduce the risk of user bias. Indeed, defining cell outlines is an important step for unconstrained TFM analysis (**Figure 1.3**) and manual contouring of hundreds of single cells is a very tedious and labor-intensive task that may introduce user-to-user variability.

To solve these problems, our algorithm identifies cell contours through a series of image-processing steps that detect both weak and strong edges despite the presence of image artifacts or fluorescent microspheres visible as black dots in bright field images. While it would also be possible to use fluorescent markers to segments cells, our aim was to keep as many fluorescence channels available as possible for other analyses and to minimize the necessary preparation steps and potential (photo-)toxicity. We therefore focused on outline detection in transmitted light images. The second algorithm speeds the statistical analysis of cell contractile output through the automated analysis of a sequence of contraction peaks. Following TFM computation, the 2-dimensional traction stress data are averaged or integrated depending on requirements for the parameters of interest within the cell area to yield a 1-dimensional time trace of contractile parameter such as total force, contraction velocity, or strain energy (**Figure 1.3**). In each trace, several contraction peaks must be analyzed to extract the relevant contractile parameters, such as contraction force amplitude, maximum contraction velocity, or maximum relaxation velocity. Our algorithm automates this task, alleviating the need for manual selection of individual peaks. Using a series of data-processing steps, including filtering in the Fourier space and auto/cross-correlations, individual peaks are automatically identified, precisely localized, extracted, aligned, and averaged in each signal trace (**Figure 1.3**). From this force-versus-time trace, the algorithm identifies the peak and baseline and computes metrics including peak amplitude, duration, mid-peak duration, maximum and minimum derivatives (for example contraction and relaxation velocities), frequency, and integral under the curve (impulse). Each metric is accompanied by its standard deviation. For example, deviation from the beating frequency is a measure of the rhythmicity of beating. We validated our code with a TFM Benchmarking Model, which we made available as a tool independent of `ConTraX`. ^64^

##### Input–Output

The inputs for *Streamlined TFM* are video recordings of fluorescent microspheres displaced by contracting hiPSC-CMs and at least one brightfield image of the cell for outlining. File formats can be *.tif*, *.avi*, or *.czi*. For *.tif* and *.avi* files, the corresponding frame rate and pixel-to-micron conversion must be provided; for *.czi* files, this information is automatically extracted from the metadata.

*Streamlined TFM* outputs a spreadsheet file containing the summary of the statistical information for the batch of cells analyzed, as well as a data folder for each cell, containing data details, cell outline mask, and images of the contraction traces.

##### Workflow

The first step after importing the input data is to verify the video parameters and to define the cell outlines. This definition can be done manually or automatically using our new cell outlining algorithm. Alternatively, if a cell outline mask was previously defined, then the user is prompted to automatically import this mask when loading the data. Cell outline masks can also be imported manually later. The user can then define a cropping mask and modify the parameters of the analysis mask (yellow dashed line in **Figure 1.2** and **Figure 1.3**), which is defined by anisotropic expansion of an ellipse fitting the cell outline. Finally, the user decides whether cropping, binning, and denoising should be performed. Each of these operations can significantly shorten the processing time and reduce noise in the output data, but should be carefully considered and reported if used, as they may introduce a consistent bias into the results depending on image quality. The *TFM Benchmarking Model* (see online Detailed Methods) is a useful tool for exploring the impact of these settings.

The second step measures the deformation induced in the hydrogel by contracting hiPSC-CMs by tracking the displacement of the fluorescent microspheres using digital image correlation (DIC). In this step, the user should verify that adequate DIC parameters are used; again, the *TFM Benchmarking Model* is helpful. After DIC computation, the user must confirm or select a correct reference frame for the contracted and relaxed state of the cell, save the data, and proceed to the next step.

The third step computes the traction stresses from the strain fields through Fourier Transform Traction Cytometry (FTTC). This step does not require specific user input, but for the choice of analysis to perform (constrained or unconstrained analysis). The regularization parameter, which is crucial for TFM computation, is calculated automatically using L-curve corner detection.^15,63^

Finally, the results panel displays the computed time traces for each parameter. While the automated peak analysis automatically analyzes each trace parameter and saves them in output files, the user can manually confirm or correct individual parameters, which are then saved in a separate tab in the spreadsheet. The user can select flags and add comments to specific cells for downstream analysis.

## ONLINE FIGURES AND LEGENDS

**Figure S1:**
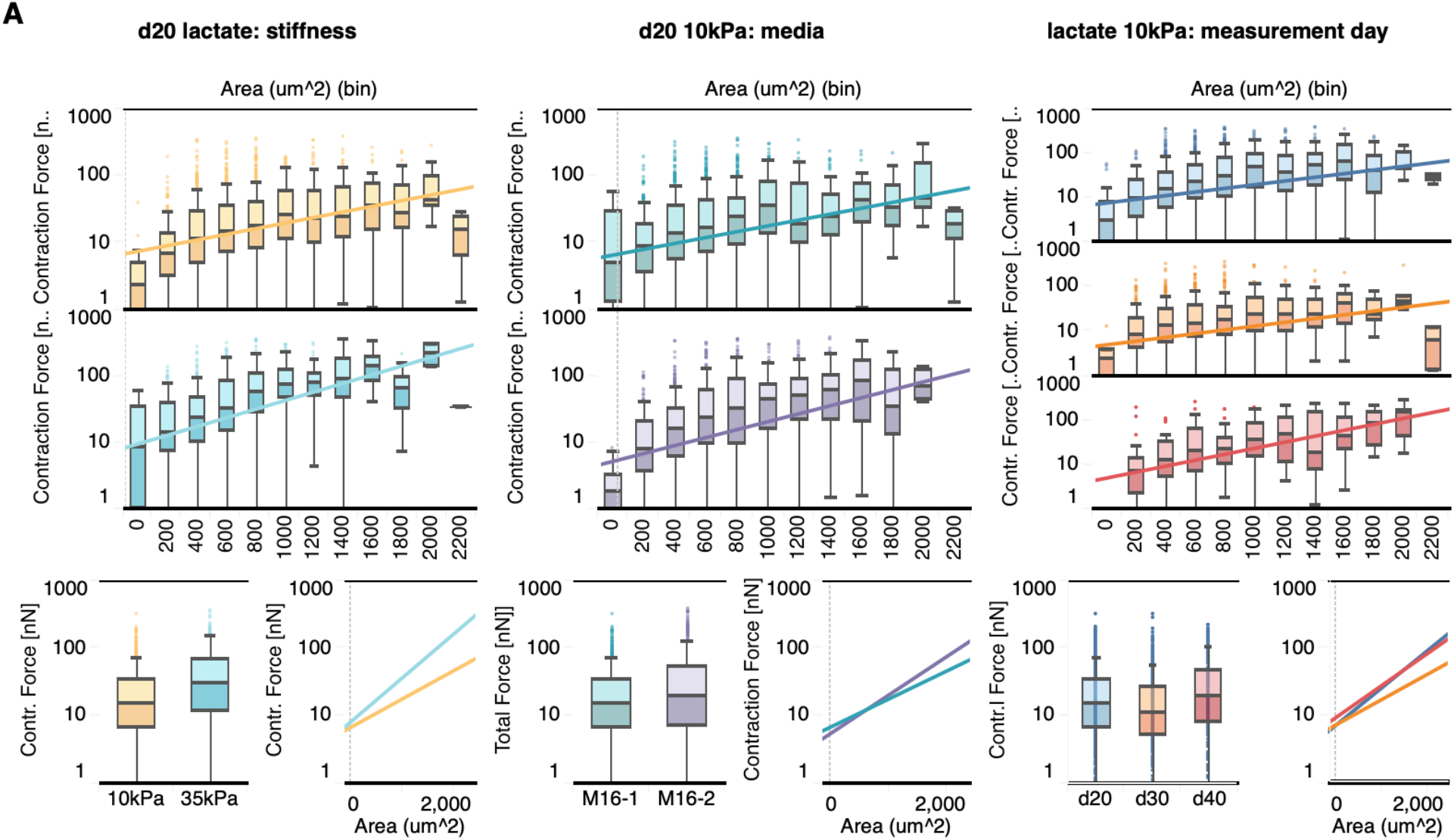
Contractile force depends on cell area and on substrate stiffness.

**Figure S2:**
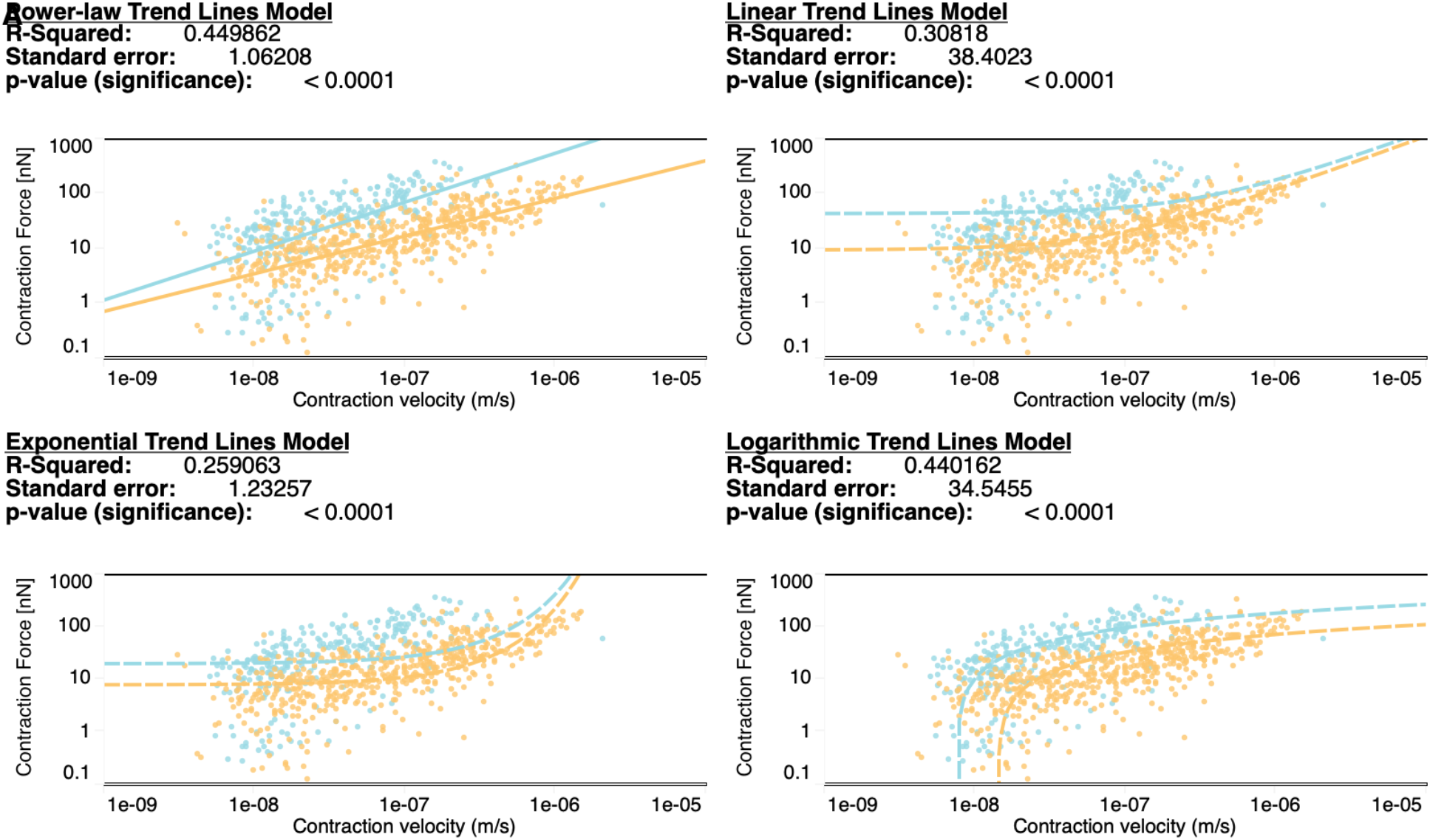
Contraction forces are impacted by substrate stiffness at day 20 in M16_lac_ medium and are best fitted by a power-law relationship.

### Power-law Trend Lines Model

A linear trend model was computed for the natural log of Contraction Force [nN] given the natural log of Contraction velocity (m/s). The model is significant at *p* <= 0.05, meaning that there is a non-zero-slope linear relationship between the natural log of Contraction Force [nN] and the natural log of Contraction velocity. The difference in model fit as a factor Stiffness is ignificant at *p* <= 0.05.

|  |  |
| --- | --- |
| **Model formula:** | Stiffness*(ln(Contraction velocity (m/s)) + intercept) |
| **Number of modeled observations:** | 837 |
| **Number of filtered observations:** | 17 |
| **Model degrees of freedom:** | 4 |
| **Residual degrees of freedom (DF):** | 833 |
| **Sum squared error (SSE):** | 939.632 |
| **Mean squared error (MSE):** | 1.12801 |
| **R-Squared:** | 0.449862 |

**Analysis of Variance:**
| <u>Field</u> | <u>DF</u> | <u>SSE</u> | <u>MSE</u> | <u>F</u> | <u>p-value</u> |
| --- | --- | --- | --- | --- | --- |
| Stiffness | 2 | 258.19928 | 129.1 | 114.449 | < 0.0001 |

**Individual trend lines:**
| <u>Stiffness</u> | <u>p-value</u> | <u>DF</u> | <u>Term</u> | <u>Value</u> | <u>SE</u> | <u>t-value</u> | <u>p-value</u> |
| --- | --- | --- | --- | --- | --- | --- | --- |
| 35kPa | < 0.0001 | 284 | ln(Contraction velocity (m/s)) | 0.891683 | 0.0686588 | 12.9872 | < 0.0001 |
|  |  |  | intercept | 18.5952 | 1.19143 | 15.6074 | < 0.0001 |
| 10kPa | < 0.0001 | 549 | ln(Contraction velocity (m/s)) | 0.686513 | 0.0308868 | 22.2267 | < 0.0001 |
|  |  |  | intercept | 13.8682 | 0.507475 | 27.3279 | < 0.0001 |

**Figure S3.**
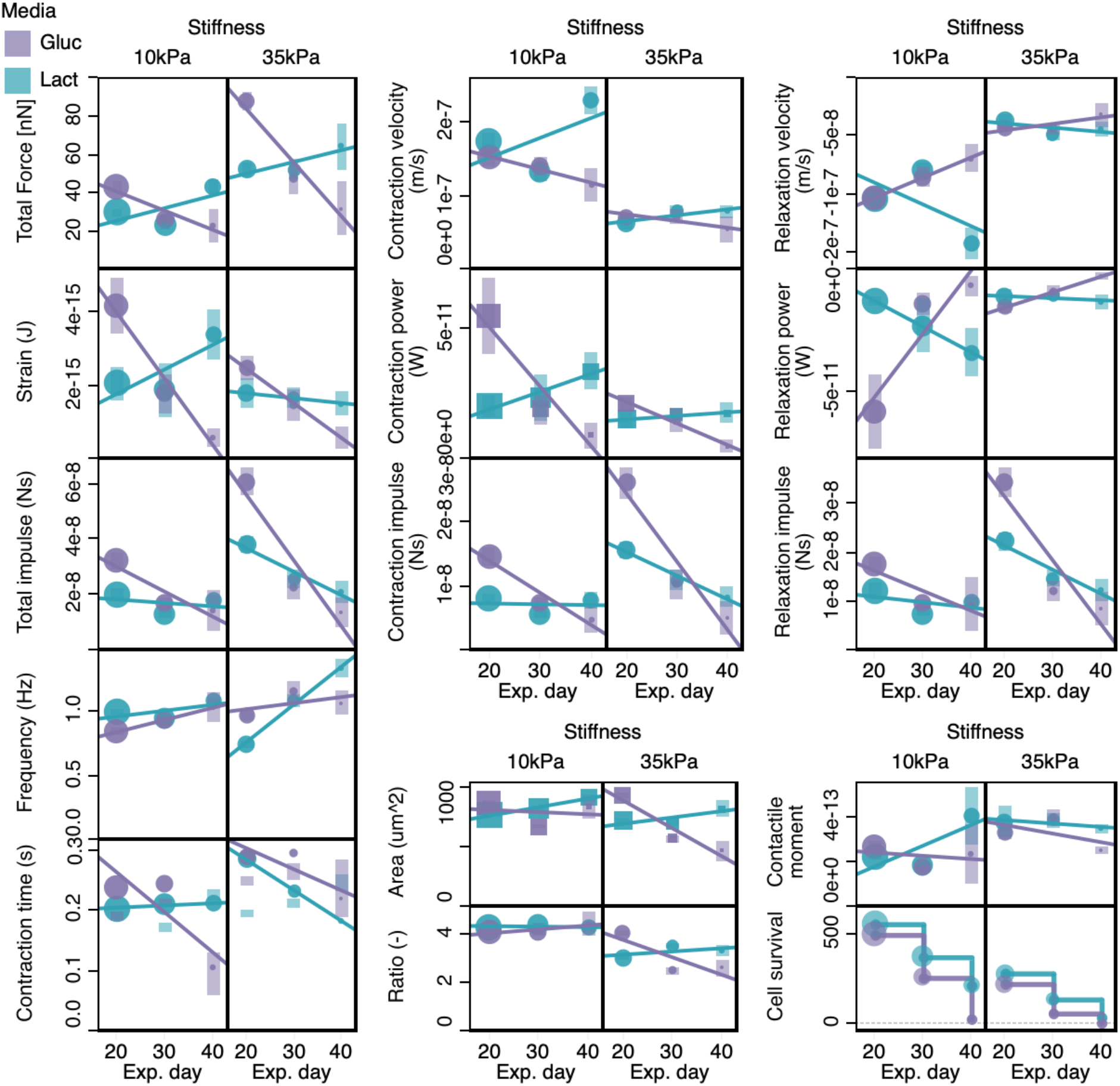
Evolution of all the measured contractile parameters over the time course experiment for both substrate stiffness and medium compositions.

**Figure S4:**
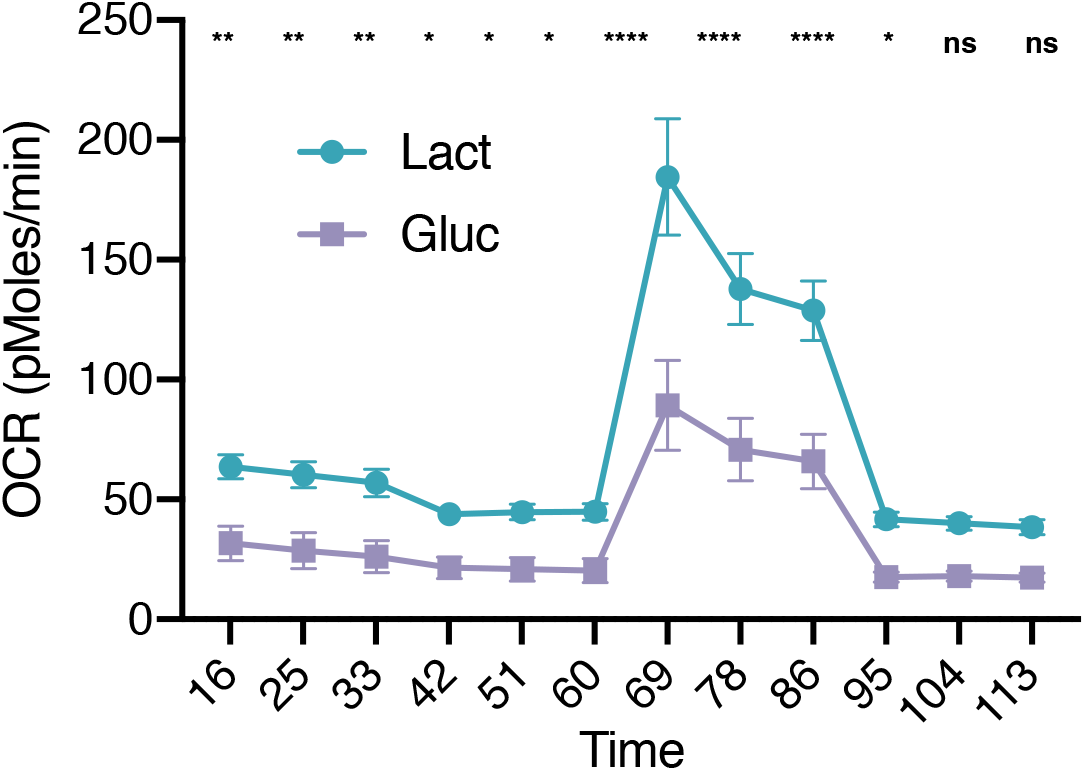
Seahorse metabolic assay for hiPSC-CMs cultured in M16_lac_ and M16_glu_ media. Mean basal and maximaloxygen consumption rate (OCR) is higher for cells in M16**lac** medium. Statistics with t-test: \**p* < 0.05, \*\**p* < 0.005, \*\*\**p* < 0.001, \*\*\*\**p* < 0.0001. ns, not significant, 3 biological replicates, >3 technical replicates.

